# Iron deficiency drives metabolic adaptation of red pulp macrophages via ferroportin-SYK signaling and BCAA catabolism to enhance erythrophagocytosis

**DOI:** 10.1101/2025.06.13.659526

**Authors:** Pratik Kumar Mandal, Raghunandan Mahadeva, Komal Chouhan, Patryk Slusarczyk, Gabriela Zurawska, Marta Niklewicz, Matylda Macias, Aleksandra Szybińska, Aneta Jończy, Zhaoyuan Liu, Florent Ginhoux, Małgorzata Lenartowicz, Wojciech Pokrzywa, Elizabeta Nemeth, Katarzyna Mleczko-Sanecka

## Abstract

Iron deficiency is the most common nutritional disorder worldwide, yet how individual cell types adjust to iron scarcity remains unclear. Splenic red pulp macrophages (RPMs) are essential for systemic iron homeostasis. They manage exceptionally high iron flux by recycling aged red blood cells through erythrophagocytosis, a specialized form of efferocytosis. However, how RPMs adapt their clearance and metabolic programs to iron deficiency remains unexplored. Here, we show that RPMs from mildly anemic, iron-deficient mice exhibit enhanced erythrophagocytic capacity. Proteomic profiling and flow cytometry revealed expansion and activation of lysosomal and mitochondrial networks, accompanied by elevated mitochondrial respiration. This metabolic rewiring and the increase in erythrophagocytosis depended on branched-chain amino acid (BCAA) catabolism. These responses were distinct from alternative macrophage activation states and absent in liver and peritoneal macrophages. Mechanistically, the low hepcidin-high ferroportin axis and SYK kinase activity emerged as drivers of this functional rewiring. Pharmacological inhibition of SYK or BCAA catabolism blunted iron-deficiency-induced erythrophagocytic and mitochondrial adaptations of RPMs. Together, these findings reveal a non-canonical metabolic reprogramming of RPMs that enhances their specialized clearance functions during systemic iron scarcity, raising possiblity that similar signaling circuits may operate in other efferocytic macrophage subsets under altered ferroportin-SYK axis activity.

## Introduction

Efficient clearance of apoptotic cells, termed efferocytosis, is fundamental to tissue homeostasis and the prevention of inflammation by averting the release of potentially damaging intracellular components into the surrounding microenvironment^1–4^. Macrophages, the primary phagocytes responsible for efferocytosis, undergo profound metabolic reprogramming to sustain this process^5^. Lipid metabolite-driven receptor activation (e.g., PPARδ, LXR) promotes continual efferocytosis^6–8^ coupled with enhanced glycolysis^9,10^, fatty acid oxidation^11^, mitochondrial respiration and dynamics^12,13^, and an upregulated pentose phosphate pathway^14,15,61^. Among amino acids, the metabolism of arginine [via arginase 1 (ARG1) and ornithine decarboxylase (ODC1)]^16^, methionine (leading to modulation of DNA methylation)^17^, tryptophan [by indoleamine 2,3-dioxygenase 1 (IDO1)]^18^, and glutamine [via glutaminase-1 (GLS1)-mediated glutaminolysis]^19^ have well-characterized roles in regulating macrophage function during efferocytosis. However, a critical gap remains in our understanding of how diverse metabolic changes are integrated and regulated across macrophage subsets and specialized efferocytic processes. In particular, very little is known about the metabolic or signaling control of erythrophagocytosis (EP), the process of clearing physiologically aged or damaged red blood cells (RBCs), which is predominantly carried out by red pulp macrophages (RPMs) in the spleen^20–22^. As RBCs constitute the body’s largest iron store, their efficient removal and subsequent iron recovery are critical for supporting erythropoiesis^21,22^. Upon engulfment, RBCs are processed within phagolysosomes, releasing heme, which is catabolized by heme oxygenase 1 (HO-1) into ferrous iron (Fe^2+^) that feeds the cellular labile iron pool (LIP). Within RPMs, iron from the LIP is either stored within ferritin or exported via ferroportin (FPN) to replenish circulating transferrin-bound iron. The iron export capacity of FPN is tightly regulated by hepcidin, a liver-derived hormone that provokes occlusion and/or degradation of FPN, thus ensuring systemic iron homeostasis^23–25^. The central role of RPMs in the maintenance of iron homeostasis is underscored by the functional attenuation in RBC clearance, accompanied by progressive splenic iron accumulation, observed in mice lacking genes essential for RPM development and during physiological aging^26–28^. Emerging evidence indicates that EP is regulated by such factors as pro-inflammatory signals^29–31^, the mechanosensitive channel PIEZO1^32^ via calcium (Ca^2+^) signaling^27,32^, and iron accumulation^27^. How these or other factors interact to fine-tune EP in response to varying physiological demands, particularly in states of altered iron bioavailability, remains a key unanswered question.

The critical nature of the RPM-driven recycling system takes on new significance in the face of the global burden of iron deficiency, the main cause of anemia, which impacts approximately 1.8 billion people (approximately 30% of the population)^33–36^. According to the 2021 Global Burden of Disease study, anemia ranks as the third leading contributor to years lived with disability worldwide, and, despite its multifactorial origins, dietary iron deficiency remains the primary cause^35,37^. Critically, the prevalence of iron deficiency itself significantly exceeds that of clinically diagnosed anemia and represents a crucial intersection between cellular iron metabolism and public health outcomes, particularly in resource-constrained developing regions^36–38^. Yet, the specific cellular vulnerabilities and the molecular mechanisms orchestrating adaptation to iron deficiency, including those operating within iron-recycling splenic RPMs, remain poorly defined.

Here, we demonstrate that RPMs in iron-deficient (ID) mice exhibit enhanced EP, and we explore how this functional adjustment is regulated by cellular signaling and metabolic rewiring. We reveal that the hepcidin-FPN axis, via regulating Ca^2+^ and spleen tyrosine kinase (SYK) signaling, enhances the EP capacity of RPMs together with their mitochondrial respiration to optimize iron recycling during iron deficiency. Furthermore, we show that RPMs uniquely rely on efficient catabolism of branched-chain amino acids (BCAAs), rich in RBCs, to enhance their EP capacity, and that this metabolic requirement remains under the control of high FPN and SYK. In sum, our findings highlight novel signaling and metabolite-driven mechanisms that shape the functional plasticity of RPMs during limited iron supplies. These insights not only deepen our understanding of RPM biology but may also inform how other macrophage populations adapt to metabolic stress or iron fluctuations in diverse physiological and pathological contexts.

## Results

### Nutritional iron deficiency increases the EP capacity of RPMs

To investigate the impact of low systemic iron availability on EP and, more broadly, the iron-recycling capacity of RPMs, we employed a murine model of nutritional iron deficiency. To this end, four-week-old female mice were subjected to either an ID diet (<5 ppm iron) or an iron-balanced (IB) control diet (200 ppm iron) for five weeks. Dietary iron restriction resulted in significantly decreased splenic (Figure 1A) and hepatic (Figure 1B) non-heme iron concentrations in the ID group compared to the IB group, confirmed by reduced iron deposition within the splenic red pulp via Perl’s staining (Figure S1A). A subtle but significant reduction in cardiac (Figure S1B) non-heme iron was also observed in the ID group, while muscle (Figure S1C) tissue iron content remained unaffected. These data suggested a tissue-specific response to dietary iron restriction, with the spleen and the liver, the organs for iron recycling and storage, being the most affected. Consistent with limited iron delivery, transferrin saturation (Figure 1C) was markedly reduced in ID mice, leading to mild microcytic hypochromic anemia (Figure 1D, S1D, and S1E), which hallmarks iron deficiency. Expectedly, ID mice displayed compensatory mechanisms manifested by splenic stress erythropoiesis, as evidenced by elevated serum erythropoietin (EPO) (Figure S1F) and increased percentage of splenic, but not bone marrow, CD71^+^/TER119^+^ progenitor cells (Figure 1E and S1G).

**Figure 1.**
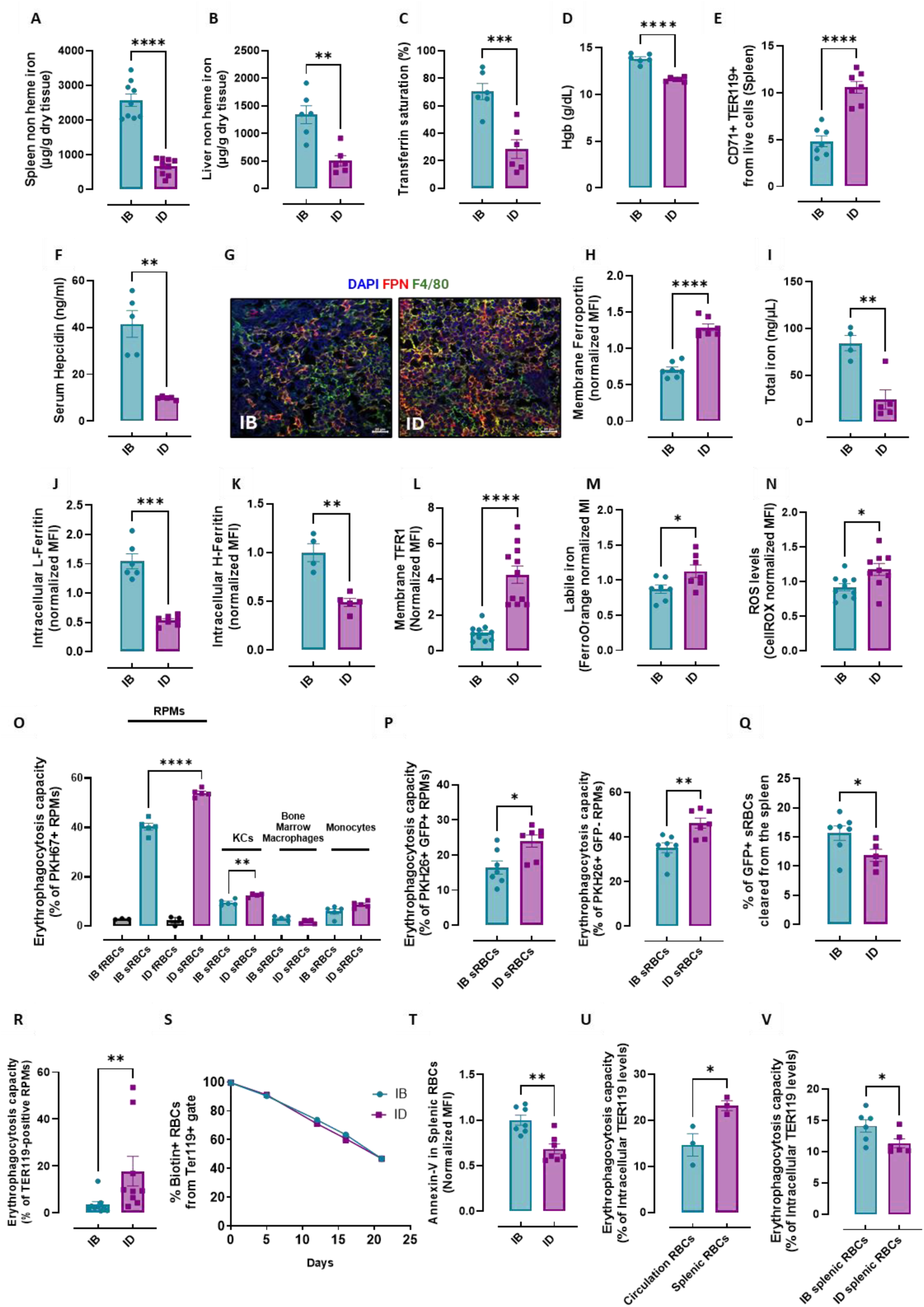
Nutritional iron deficiency increases the erythrophagocytic capacity of RPMs. (A) Spleen and (B) liver non-heme iron levels, (C) plasma transferrin saturation, and (D) blood hemoglobin (Hgb) were measured in iron-balanced (IB) and iron-deficient (ID) mice. (E) Shown is the percentage of erythroid progenitor cells (TER119^+^ CD71^+^) present in the spleen of IB and ID mice. (F) Serum hepcidin levels were measured in IB and ID mice with the Hepcidin-Murine Compete ELISA Kit. (G) Representative confocal microscopy images of ferroportin (FPN) protein localization in the splenic red pulp of IB and ID mice. Frozen spleen slices were processed and stained for RPMs (F4/80, green), FPN (red), and nuclei (blue). (H) Expression of FPN on the cell membrane of RPMs derived from IB and ID mice, gated as F4/80^high^, CD11b^dim^, was assessed by flow cytometry. (I) The total intracellular iron content in magnetically sorted RPMs from IB and ID mice was assessed using the Iron Assay Kit. (J) Intracellular L-ferritin and (K) H-ferritin protein levels in RPMs of IB and ID mice were quantified by flow cytometry. (L) Expression of TFR1 on the cell membrane of RPMs derived from IB and ID mice was assessed by flow cytometry. (M) Cytosolic ferrous iron (Fe^2+^) levels in RPMs of IB and ID mice were measured using the FerroOrange probe with flow cytometry. (N) The cytosolic ROS levels in RPMs of IB and ID mice were assessed by determining CellROX Deep Red fluorescence intensity with flow cytometry. (O) The erythrophagocytosis (EP) capacity in RPMs and other myeloid populations of IB and ID mice was determined using flow cytometry by measuring the percentage of cells that phagocytosed transfused PKH67-labeled RBCs. fRBCs indicate fresh RBCs, sRBCs - stressed RBCs (shown only for RPMs; in other indicated populations, fRBCs uptake was negligibly low). (P) IB and ID mice were transfused with GFP-expressing and PKH26-stained sRBCs. The percentage of RPMs positive for intact RBCs (PKH26^+^GFP^+^) and RBC ghosts (PKH26^+^GFP^-^) was determined. (Q) Flow cytometric analysis of splenic GFP-sRBC percentage in ID vs IB mice. (R) The endogenous EP capacity of RPMs derived from IB and ID mice was determined by intracellular staining of the RBC marker TER119 in RPMs and flow cytometry. (S) The RBC biotinylation lifespan assay was performed on circulating RBCs from IB and ID mice. (T) Flow cytometry assessment of phosphatidylserine exposure (Annexin V staining) in splenic RBCs of ID vs IB mice. (U) EP capacity by primary cultured macrophages (iRPM) was determined using splenic and circulating RBCs of IB mice. (V) EP capacity of iRPMs exposed to RBCs derived from the spleens of IB and ID mice. Each dot represents one mouse or an independent cell-based experiment. Each dot in the graph (S) represents n=9. Data are represented as mean ± SEM. Welch’s unpaired t-test determined statistical significance between the two groups. ns p>0.05, *p<0.05, **p<0.01, ***p<0.001 and ****p<0.0001.

Next, we aimed to precisely characterize cell-intrinsic iron parameters of RPMs in response to iron-restricted feeding. Consistent with the drop in serum levels of the iron-regulatory hormone hepcidin (Figure 1F), FPN expression in RPMs was significantly elevated in ID mice, as shown by F4/80 immunofluorescence in splenic red pulp and by flow cytometric analysis of F4/80^high^ CD11b^dim^ RPMs^27^ (Figure 1G and 1H). In agreement with boosted iron-export capacity, a marked decrease in total iron content was observed in the magnetically sorted RPMs from ID mice (Figure 1I), along with reduced protein expression of both L- and H-ferritin as assessed by flow cytometry (FTL and FTH1; Figure 1J and 1K), indicating depletion of intracellular iron stores. Furthermore, we detected upregulated levels of transferrin receptor 1 (TFR1) surface expression (Figure 1L), implying a cellular response to iron deficiency. However, paradoxically, despite enhanced iron export, RPMs in ID mice unexpectedly displayed a mild but significant elevation of LIP, as assessed using the Fe^2+^-specific fluorescent probe FerroOrange (Figure 1M). Consistent with this finding, we also detected mildly elevated reactive oxygen species (ROS) levels in RPMs of ID versus IB mice (Figure 1N).

Intrigued by the increased LIP in RPMs of ID mice, we investigated the capacity of EP, the major route for RPM iron acquisition. To this end, following well-established procedures^27,39^, we transfused mice with PKH67-labeled RBCs. Strikingly, flow cytometry analysis revealed significantly enhanced phagocytosis of temperature-stressed RBCs (sRBC) by RPMs from ID compared to IB mice, a phenomenon not observed for non-stressed fresh RBCs that were engulfed at comparably negligible levels (Figure 1O). In addition, we evaluated EP capacity across other myeloid populations, including Kupffer cells (KCs) in the liver, macrophages in the bone marrow, and circulating monocytes (Figure 1O). Among these, only KCs showed a statistically significant, albeit modest, increase in EP capacity in iron-restricted mice. Previous work has shown that splenic red pulp sinuses trap aged RBCs, leading to partial hemolysis and subsequent phagocytosis of membrane remnants (RBC ghosts) by RPMs^40^. To distinguish between the clearance of intact sRBCs and RBC ghosts in our model, we intravenously injected mice with sRBCs ubiquitously expressing GFP and stained with PKH26 membrane dye, as described previously^41^. Consistent with our prior results, RPMs from ID mice exhibited increased EP of GFP^+^ PKH26^+^ and GFP-PKH26^+^ sRBCs compared to control IB animals, confirming their enhanced capability for the uptake of both intact RBCs and RBC ghosts (Figure 1P). Likewise, splenic clearance of transfused GFP-positive sRBCs was accelerated in ID mice compared to IB controls (Figure 1Q), further supporting enhanced sRBC removal by RPMs within the low-iron splenic microenvironment. To validate sRBC transfusion assay results, we next assessed intracellular levels of the erythrocyte marker TER119 in RPMs as a measure of endogenous EP capacity^27,42^, confirming its elevation in ID mice (Figure 1R). Finally, we verified whether increased EP in low-iron mice is solely driven by macrophage-intrinsic mechanisms or could result from erythrocyte-derived signals. RBC lifespan remained comparable between IB and ID mice (Figure 1S), suggesting that iron restriction did not compromise RBC integrity. Next, exposure of the apoptotic cell marker phosphatidylserine in splenic RBCs was not enhanced in ID compared to IB mice (Figure 1T). Finally, using a co-culture assay of magnetically isolated splenic and circulating RBCs with primary macrophages, we found that while aged splenic RBCs are preferentially sequestered compared to circulating RBCs (Figure 1U), iron deficiency does not increase splenic RBC susceptibility to phagocytosis (Figure 1V), implying no induction of RBC “eat-me” signals by iron restriction. Taken together, we uncovered that systemic iron restriction potentiates the cell-intrinsic RBC clearance capacity of RPMs, likely aimed at balancing the increased replenishment of the plasma iron pool by FPN-mediated export.

### Iron deficiency drives metabolic reprogramming and enhanced lysosomal activity of RPMs via the non-canonical type of polarization

To decipher the molecular mechanisms of enhanced EP capacity in RPMs of ID mice and explore potential accompanying functional changes, we first investigated the transcriptome of FACS-isolated RPMs. Surprisingly, only 145 genes were differentially expressed between RPMs derived from ID and IB mice (Figure 2A, Table S1). Bioinformatic analysis revealed significant enrichment of cell cycle regulators (e.g., *Mki67, Plk1, Plk2, Aurka, E2f7, Sgo1, Kif4, Cenpf, Bub1, Ccnb1, Ccnb2*) among genes upregulated by iron deficiency. This suggested increased pro-mitotic signaling in RPM of ID mice, which was corroborated by elevated protein levels of the proliferation marker Ki67 (Figure 2B). However, this response was not reflected by any significant change in the representation of total RPMs within CD45^+^ splenic cells (Figure 2C). Likewise, using the Ms4a3Cre-RosaTdT reporter mice^43^, we found no significant differences in the relative representation of residual embryonically-derived RPMs within the RPM population (Figure 2D), suggesting the absence of their proliferation. The percentage of monocyte-derived RPMs also remained constant at approximately 15% in both IB and ID mice (Figure 2D), implying that the EP enhancement was not driven by *de novo* monocyte recruitment.

**Figure 2.**
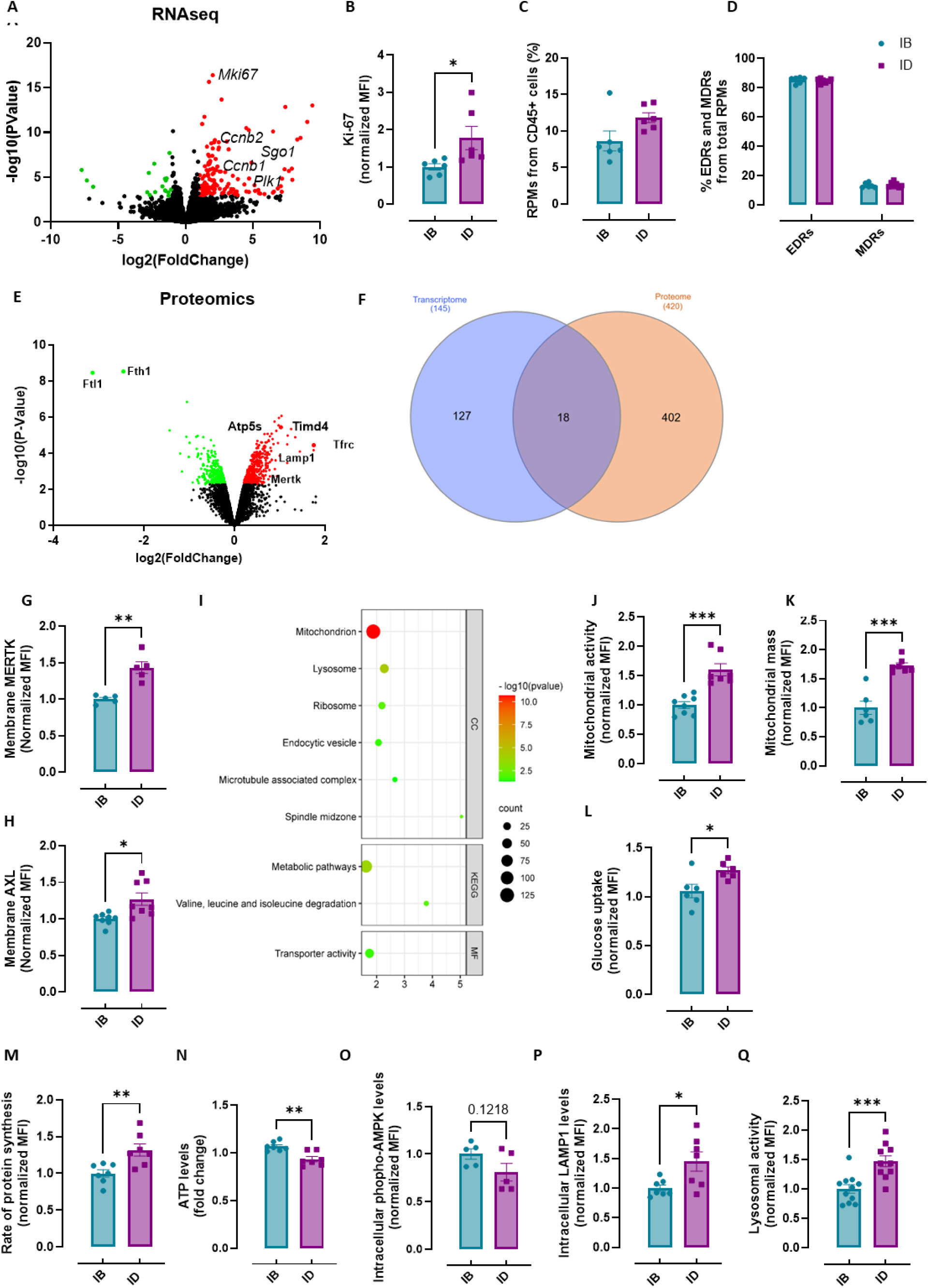
Iron deficiency drives metabolic reprogramming and enhanced lysosomal activity of RPMs. (A) A volcano plot of differentially regulated genes identified by the RNA-Seq transcriptomic analysis in FACS-sorted RPMs derived from IB and ID mice. (B) Intracellular Ki-67 protein levels in RPMs derived from IB and ID mice were assessed by flow cytometry. (C and D) Percentage of (C) total RPMs and (D) embryonic-derived RPMs (EDRs) and monocyte-derived RPMs (MDRs) among total RPMs in IB and ID mice. (E) Volcano plot visualizing the proteomic landscape of RPMs in response to iron deficiency, depicting differentially expressed proteins in IB and ID mice. (F) Venn diagram comparing the differentially expressed genes (transcriptome) and proteins (proteome) in RPMs from ID mice. (G and H) Cell membrane expression levels of (G) MERTK and (H) AXL of RPMs in IB and ID mice were determined by using fluorescently labeled antibodies and flow cytometry. (I) Enriched functional categories among significantly increased protein hits in magnetically sorted RPMs derived from ID versus IB spleen. (J) Mitochondrial membrane potential/activity and (K) mitochondrial mass were determined in RPMs derived from IB and ID mice using TMRE probe and MitoTracker Green, respectively, with flow cytometry. (L) Quantification of glucose uptake via flow cytometry in RPMs from IB and ID mice using a fluorescent glucose analog. (M) Flow cytometric detection of newly synthesized proteins in RPMs of IB and ID mice using Click-iT Plus OPP Alexa Fluor 488 Protein Synthesis Assay. (N) Quantification of ATP levels in FACS-sorted RPMs (10,000 cells) from IB and ID mice using a chemiluminescence-based assay. (O) Intracellular phosphorylated AMPK protein levels in RPMs derived from IB and ID mice were assessed by flow cytometry. (P) Intracellular expression levels of the core lysosomal protein LAMP1 in RPMs of IB and ID mice were determined by flow cytometry. (Q) Flow cytometric assessment of lysosomal activity in RPMs of IB and ID mice using the Lysosomal Intracellular Activity Assay Kit. Each dot represents one mouse. Data are represented as mean ± SEM. Welch’s unpaired t-test determined statistical significance between the two groups. ns p>0.05, *p<0.05, **p<0.01, ***p<0.001 and ****p<0.0001.

To further explore the adaptation of RPMs to systemic iron deficiency, we employed TMT-based quantitative proteomics of magnetically-sorted F4/80-high RPMs. This analysis identified 420 proteins significantly more abundant and 282 decreased in RPMs of ID mice compared to IB mice (Figure 2E, Table S2). Confirming the quality of the proteomics analysis, we found FTH1 and FTL as the top downregulated hits, and TFR1 as the most induced protein in RPMs of iron-restricted mice. Consistent with this expression pattern, IRP2, a regulator of cellular iron balance, was stabilized in RPMs of ID mice. In agreement with RNA-seq data, we also detected an increase in several regulators of cell cycle progression, such as AURKA, AURKB, PLK1, and CDK1 in RPMs of low-iron mice. Intriguingly, proteomic analysis revealed a pronounced functional rewiring of RPMs in ID mice, highlighting significant post-transcriptional regulation, with far more protein-level alterations and minimal overlap between transcriptomic and proteomic datasets (Figure 2F). First, we observed upregulation of the phosphatidylserine-recognizing phagocytic receptors MERTK and TIM4 (significant hits), and a trend towards induction of AXL and STAB2, along with the Fc receptor CD16 in RPMs of ID mice, responses that were all confirmed by flow cytometry (Figure 2G-H, Figure S2.1A-C). Interestingly, this elevated capacity for recognizing “eat me” signals was accompanied by the induction of the “don’t eat me” receptor SIRPα on RPMs in ID mice (Figure S2.1D), hinting at a mechanism that promotes the clearance of aged dysfunctional RBCs, sparing those that are intact. Next, to gain unbiased insights into processes supporting the enhanced EP capacity of RPMs in ID mice, we performed functional enrichment analyses on the differentially expressed proteins. Notably, mitochondrial proteins emerged as the most significantly enriched category among hits upregulated in RPMs of ID compared to IB mice (Figure 2I), including factors involved in ATP synthesis (ATP5S), metabolite transport (SLC25A5, VDAC1), mitochondrial protein import (SAMM50), iron-sulfur cluster assembly (SFXN2), cristae remodeling (MCCC1, PMPCB), and coenzyme Q-10 biosynthesis (COQ9, COQ5). Flow cytometric analysis confirmed these expanded mitochondrial functions by demonstrating a significant increase in both mitochondrial membrane potential (a proxy for mitochondrial activity) (Figure 2J) and mitochondrial mass (Figure 2K) in RPMs of ID mice. Consistently, RPMs of ID *versus* IB mice exhibited increased uptake of fluorescently labeled glucose analog (Figure 2L) and showed a significant upregulation of protein synthesis (Figure 2M), a major ATP-consuming process reflective of enhanced cellular bioenergetics. Interestingly, ATP levels in FACS-sorted RPMs revealed a mild depletion in ID compared to IB mice (Figure 2N), but the activation status of AMPK, a key sensor of cellular energy stress, was not enhanced (Figure 2O). This suggests that RPMs of ID mice did not exhibit energy deficit, but rather a state where increased ATP production is insufficient to meet the high ATP demand driven by accelerated EP and protein synthesis. Importantly, in contrast to RPMs, neither peritoneal macrophages (not involved in iron recycling) nor iron-recycling KCs show hallmarks of metabolic rewiring in ID mice, as exemplified by unaltered mitochondrial activity and mass (Figure S2.1E-H).

Further analysis of the proteomic data revealed another major category, encompassing proteins associated with lysosomal functions, as enriched in RPMs of ID compared to IB mice (Figure 2I). In total, RPMs of iron-restricted mice increased the protein levels of 25 key components of the Coordinated Lysosomal Expression and Regulation (CLEAR) network, a transcriptional program that drives lysosomal biogenesis and function^44^ (Figure S2.2A). This included integral lysosomal membrane proteins, LAMP1 and LAMP2, vacuolar ATPase subunits (ATP6V0D1, ATP6V0A1), essential for lysosomal acidification, and a diverse array of lysosomal hydrolases, including cathepsins B, D, and S; α-glucosidase (GAA); β-glucosidase (GBA); N-acetylglucosaminidase (NAGA); and sialidase 1 (NEU1). Flow cytometric analysis confirmed elevated intracellular LAMP1 levels (Figure 2P), indicating increased lysosomal mass in RPMs of ID mice. Consistent with this, we observed enhanced lysosomal activity in RPMs of ID mice using a fluorescent probe (Figure 2R). Further ultrastructural analysis of RPMs within splenic sections with TEM revealed that while the number of regular lysosomes, autophagolysosomes, autophagosomes, and peroxisomes remained unaltered between control and low-iron mice, large lysosomes were more abundant in RPMs of iron-restricted mice (Figure S2.2B and S2.2C). While lysosomal size is dynamic and influenced by nutrient availability, emerging evidence, for instance, from senescent cells suggests that a higher proportion of enlarged lysosomes correlates with increased proteolytic activity and lysosomal mass^45,46^, remaining consistent with our observations. Taken together, these data suggest that enhanced lysosomal function in RPMs of ID mice is coordinated with heightened RBC clearance, extending the spectrum of RPMs reprogramming in iron deficiency.

Erythrocytic heme and iron loading, as well as systemic or cell-intrinsic iron deficiency, have been shown to influence macrophage polarization, promoting either pro-inflammatory or anti-inflammatory profiles, though findings have often been contradictory^47–50^. Since metabolic reprogramming is closely linked to macrophage polarization, which in turn shapes the outcome of infections and inflammatory diseases^51^, we assessed the polarization state of RPMs from iron-restricted mice. Our RNA-seq and proteomics data revealed no clear induction of either pro- or anti-inflammatory mediators, alongside a mixed expression pattern of surface signals identified by flow cytometry. While some markers typically associated with M1 (MHCII, CD74) and M2 (CD206) polarization were decreased in RPMs of ID *versus* IB mice, others, such as the pro-inflammatory CD38 and the anti-inflammatory CD163, were increased (Figure S2.3). This atypical combination of features suggests that augmented RBC clearance in nutritionally iron-restricted mice leads to the activation state of RPMs distinct from the canonical M1 or M2 type of profile.

### Low hepcidin levels are critical for functional adaptations of RPMs during iron deficiency

To elucidate the mechanisms driving the enhanced EP capacity of RPMs in iron deficiency, we first investigated whether these adaptations were directly dependent on intracellular iron levels. Iron depletion *in vitro* using deferiprone (Figure 3A) or DFO^27^ did not alter the RBC clearance capacity of primary BMDM macrophages cultured with hemin (iRPM)^27^. Furthermore, our *in vivo* data demonstrated that despite systemic iron deficiency, RPMs in ID mice exhibited elevated LIP (Figure 1M), likely due to their increased EP activity, indicating that iron chelation *in vitro* might not fully recapitulate the complex cellular interactions and signaling pathways present in RPMs *in vivo*. These findings suggested that factors beyond simple iron availability may be critical drivers of enhanced RPM phagocytic activity during iron deficiency. To address this possibility mechanistically, we developed an *in vitro* system using iRPMs cultured in serum derived from IB and ID mice, instead of bovine serum. This system recapitulated increased FPN and TFR1 expressions in iRPMs exposed to ID serum, without a significant change in total iron levels (Figure 3B-D). Notably, cells cultured in ID serum exhibited enhanced EP (Figure S3A), suggesting that serum-borne factor is sufficient to rewire the phagocytic capacity of iRPMs towards RBCs. We first excluded the contribution of EPO, the cytokine previously implicated as a stimulator of efferocytosis^52^. We showed that iRPMs, in contrast to BMDMs cultured without hemin, which is critical to mimic the heme and iron-rich nature of RPMs, did not enhance EP even under high EPO doses (Figure S3B-C). Moreover, PPARγ, a factor proposed to act downstream of EPO^52^, failed to alter the EP capacity of iRPMs (Figure S3C). Next, inspired by a study that revealed low hepcidin, rather than iron deficiency itself, as a key driver of pro-inflammatory responses in iron-restricted mice^50^, we hypothesized that reduced serum hepcidin levels and enhanced FPN stabilization in RPMs might play a critical role in the adaptive reprogramming of these cells. Strikingly, a 4-h treatment with the hepcidin mimetic PR73 reversed the enhanced EP phenotype in iRPMs exposed to ID, but not to IB serum (Figure 3E and S3D). Likewise, we conducted live imaging of EP by cultured iRPMs incubated with serum from IB and ID mice, and exposed to sRBCs labeled with a dedicated pHrodo™ dye that remains non-fluorescent at neutral pH but emits fluorescence in acidic environments, indicating sequestrations in phagolysosomes. In line with our flow cytometry results, we observed enhanced pHrodo signal in real time in iRPMs treated with ID serum, reflecting their enhanced EP capacity (Figure 3F). This boost of RBC clearance capacity was significantly abrogated upon 4 h of PR73 treatment (Figure 3F). Consistent with our *in vivo* observations, ID serum induced a significant increase in both mitochondrial and lysosomal activity, which was likewise abrogated by PR73 treatment (Figure S3E-H).

**Figure 3.**
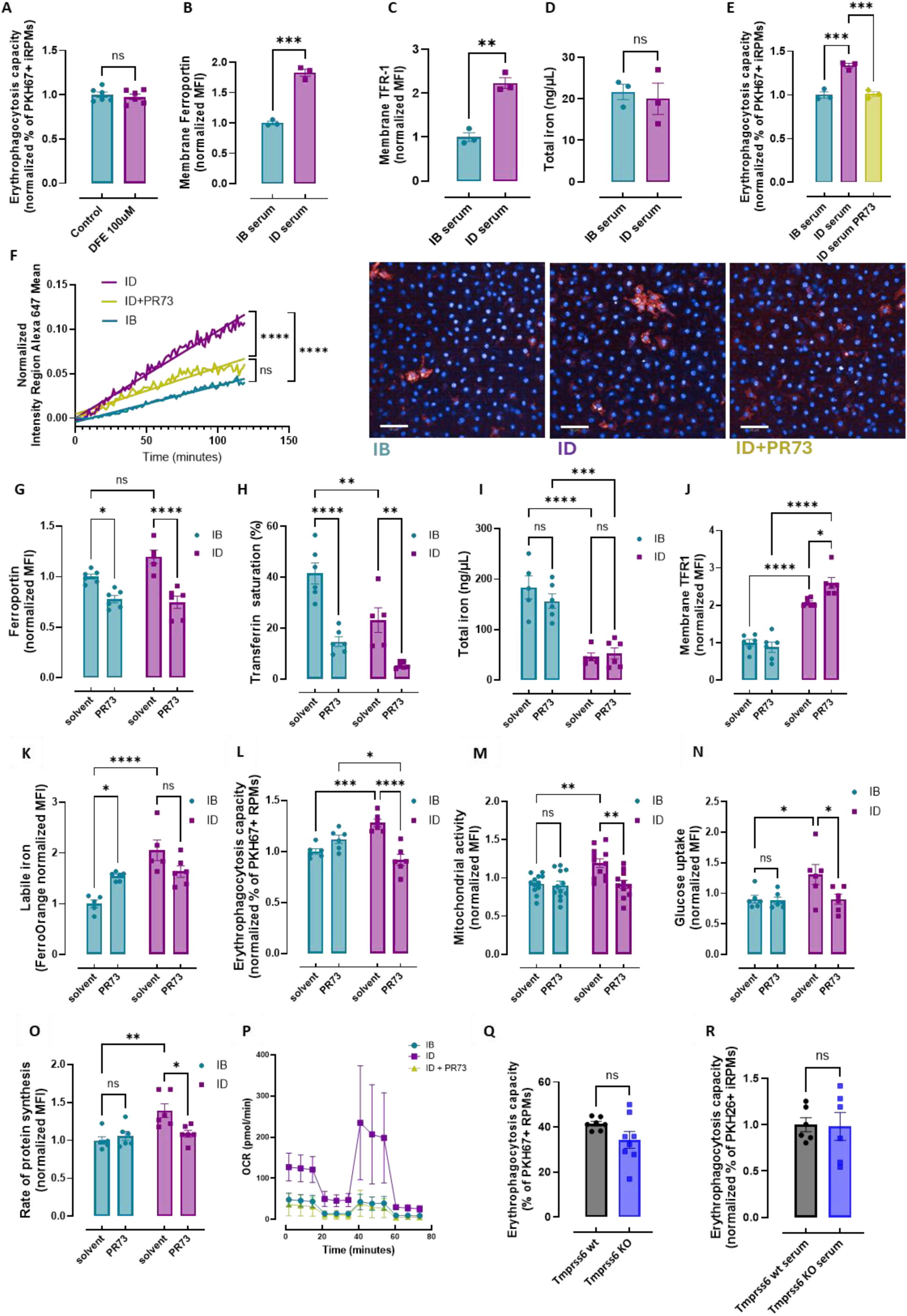
Low hepcidin levels are critical for functional adaptations of RPMs during iron deficiency. (A) Normalized EP capacity of PKH67-stained sRBCs by cultured iRPMs treated with an iron chelator DFE (100 µM, 18 h). (B-C) Flow cytometry analysis of (B) surface FPN and (C) TFR1 expression on iRPMs cultured with serum from IB or ID mice. (D) The total intracellular iron content in iRPMs cultured with serum from IB or ID mice was assessed using the Iron Assay Kit. (E) The EP capacity in iRPMs cultured with serum from IB or ID mice, and serum from ID mice supplemented with the hepcidin agonist PR73 (2 µg/mL, 4 h) determined using flow cytometry by measuring the percentage of cells that phagocytosed PKH67-labeled RBCs. (F) Time-lapse imaging of the EP progression in iRPMs, exposed to pHrodo-stained sRBCs and cultured with the serum from IB and ID mice (with and without 4 h pretreatment with PR73). The left panel shows the quantification of the RBC fluorescent signal over time, normalized to the number of macrophages. The right panel shows example images at the endpoint of the assay. Scale bar=50 μm. (G) Flow cytometric analysis of FPN cell surface expression on RPMs from IB and ID mice following intraperitoneal administration of the hepcidin agonist PR73 (50 nmol/mouse, 4 h). (H) Plasma transferrin saturation was determined in IB and ID mice following PR73 administration. (I) Intracellular iron content in magnetically sorted RPMs from IB and ID mice post-PR73 treatment as determined using the Iron Assay Kit. (J) Flow cytometric evaluation of TFR1 surface expression on RPMs derived from IB and ID mice post-PR73 administration. (K) Flow cytometric measurement of cytosolic ferrous iron (Fe^2+^) levels in RPMs from IB and ID mice using FerroOrange following PR73 administration. (L) Normalized EP capacity of PKH67-sRBCs by RPMs from IB and ID mice after PR73 administration. (M) Mitochondrial activity was determined in RPMs derived from IB and ID mice using the TMRE probe post-PR73 administration, with flow cytometry. (N) Quantification of glucose uptake via flow cytometry in RPMs from IB and ID mice upon PR73 administration using a fluorescent glucose analog. (O) Flow cytometric detection of newly synthesized proteins in RPMs of IB and ID mice using Click-iT Plus OPP Alexa Fluor 488 Protein Synthesis Assay after PR73 administration. (P) Seahorse analysis of *ex vivo* RPMs from IB and ID mice, with or without 4 h PR73 preincubation for RPMs from ID mice. (Q) Flow cytometric assessment of EP capacity in RPMs of *Tmprss6* wt and *Tmprss6* KO mice was quantified as the percentage of cells engulfing PKH67-labeled sRBCs. (R) Normalized EP capacity of PKH26 sRBCs by cultured iRPMs treated with serum from *Tmprss6* wt and *Tmprss6* KO mice was determined using flow cytometry. Each dot represents one mouse or an independent cell-based experiment. Data are represented as mean ± SEM. Welch’s unpaired t-test determined statistical significance in A-D and Q-R, while two-way ANOVA with Tukey’s Multiple Comparison tests was used in G-O. A one-way ANOVA test with Dunnett’s or Tukey’s Multiple Comparison tests was used in E. In F, the significant differences between the slopes of simple linear regression lines are shown. ns p>0.05, *p<0.05, **p<0.01, ***p<0.001 and ****p<0.0001.

To follow these *in vitro* observations, we next administered IB and ID mice with PR73 via intraperitoneal injection 4 h before analysis. PR73 administration effectively reduced cell-surface FPN expression and induced hypoferremia in both dietary groups (Figure 3G-H). However, total RPM iron content remained unchanged in both IB and ID mice following PR73 treatment (Figure 3I). PR73 administration did not reverse the increased cell surface expression of TFR1 on RPMs of ID mice but instead led to its further elevation, in contrast to IB mice, where TFR1 levels remained unaltered (Figure 3J). Interestingly, PR73 elicited similarly distinct effects on the LIP in RPMs depending on the dietary regimen: it expectedly increased LIP levels in IB mice, but led to a mild decrease in ID mice (Figure 3K). The latter pattern was, strikingly, closely reflected by the pronounced reversal of the elevated EP capacity of RPMs from ID mice by PR73 injection (Figure 3L). Likewise, PR73 abrogated most of the other hallmarks of RPM adaptation to iron deficiency, including elevated mitochondrial activity, increased glucose uptake, and heightened protein synthesis (Figure 3M-O). Intriguingly, these modulating effects of PR73 were absent in IB mice. Furthermore, *ex vivo* freshly isolated RPMs exhibited substantially elevated oxidative phosphorylation (OXPHOS), as assessed by oxygen consumption rate measurement with Seahorse (Figure 3P). Strikingly, a 4-h preincubation of the *ex vivo* RPMs of ID mice with PR73 completely abolished their activated oxygen consumption (Figure 3P). Of note, these rapid effects of PR73 did not include changes in the mitochondrial mass of RPMs, remaining elevated regardless of PR73 injection in ID mice (Figure S3I). Similarly, lysosomal activity of RPMs of ID mice remained higher than in IB mice despite PR73 supplementation, suggesting either distinct regulatory mechanisms or differential kinetics governing these responses (Figure S3J). To support the key role of the hepcidin-FPN axis in modulating the iron-recycling activity of RPMs, we employed *Tmprss6* knockout (KO) mice, a model of iron-refractory iron deficiency anemia (IRIDA)^53–55^. This model exhibits some characteristics of the nutritional ID model, including reduced transferrin saturation, decreased hepatic iron stores (Figure S3K-L), and anemia. However, in contrast to dietary ID conditions, *Tmprss6* KO mice display elevated hepcidin levels, and their splenic iron stores remain largely unchanged^56,57^, with no significant depletion observed, consistent with our findings (Figure S3M). Importantly, we observed no significant difference in RPMs’ EP capacity between *Tmprss6* KO mice and wild-type controls (Figure 3Q). Likewise, *in vitro* cultured iRPMs showed no difference in RBC clearance when exposed to serum from either *Tmprss6* KO or wild-type mice (Figure 3R). In sum, these data implied that deficient hepcidin levels and FPN stabilization, hallmarks of nutritional ID, are key driving factors for enhanced RPM erythrophagocytic capacity and their metabolic adaptation to low systemic iron levels.

### Elevated FPN regulates RPM rewiring in ID conditions via Ca²-dependent signaling

To further assess the role of FPN upregulation in RPM rewiring under systemic ID conditions, we used lentiviral transduction to create iRPMs with enforced FPN overexpression (FPN OE; FPN-iRPMs) and assessed whether this mimics RPM adaptation to iron deficiency. Control iRPMs were generated using an empty vector without the *Slc40a1* gene, encoding FPN (mock-iRPMs). We confirmed significant FPN elevation on the surface of FPN-iRPMs compared to control cells (Figure 4A). This further led to the elevation of TFR1 surface expression but did not significantly change the total iron levels (Figure 4B-C), thus resembling the iRPMs exposed to ID vs IB serum. Under basal conditions, we quantified the LIP in both FPN-iRPMs and mock-iRPMs, finding no significant difference (Figure 4D). In an attempt to recapitulate the mildly elevated LIP observed in RPMs of ID mice during enhanced *in vivo* EP, we next sought to mimic the splenic microenvironment by continuously exposing iRPMs to sRBCs during culture. Strikingly, sRBC-exposed FPN-iRPMs exhibited a significant increase in LIP compared to mock-iRPMs (Figure 4D), mirroring the *in vivo* RPM response (Figure 1M). Most importantly, FPN-iRPMs displayed enhanced RBC clearance regardless of sRBC exposure during culture (Figure 4E), a response that was efficiently reversed by suppression of FPN by a potent specific inhibitor vamifeport^58^ (Figure 4F). These findings demonstrate that forced FPN expression and consequent increased cellular iron flux, rather than overall iron content, specifically stimulate EP.

**Figure 4.**
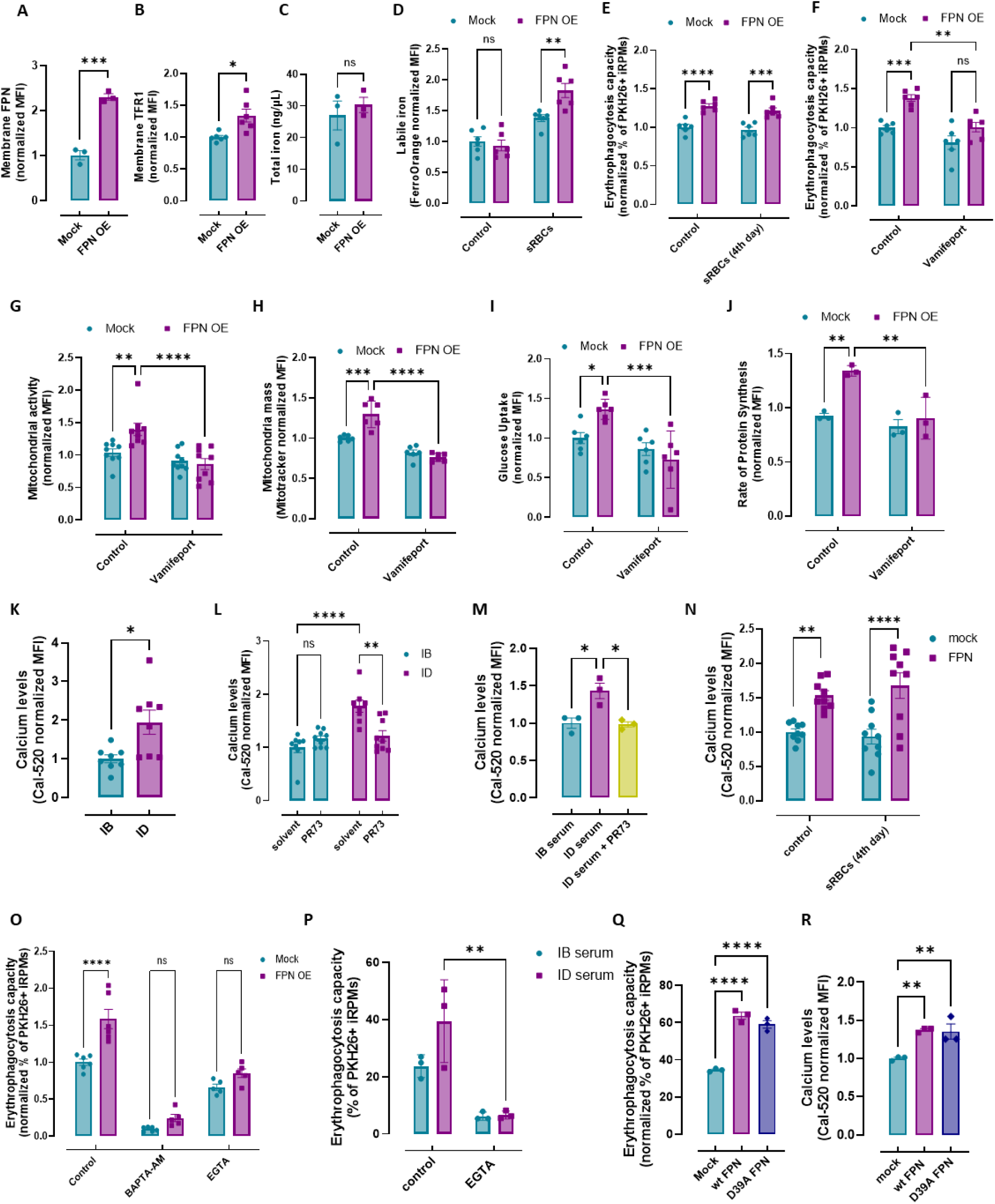
Elevated FPN regulates RPM rewiring in ID conditions via Ca²-dependent signaling. (A-J) Lentiviral transduction was used to generate iRPMs overexpressing FPN (FPN OE; FPN-iRPMs) or expressing an empty vector (mock). (A-C) Flow cytometric analysis of (A) FPN and (B) TFR1 cell surface expression, and (C) intracellular iron content in mock and FPN-iRPMs determined using the Iron Assay Kit. (D) Flow cytometric measurement of cytosolic ferrous iron (Fe^2+^) levels in mock and FPN-iRPMs with and without sRBC exposure during culture. (E) Normalized EP capacity towards PKH26-sRBCs by mock and FPN-iRPMs with and without sRBC exposure during culture was determined with flow cytometry. (F-J) iRPMs were cultured with sRBCs and treated with vamifeport (20 μM, 15 h). (F) Normalized EP capacity towards PKH26-sRBCs was determined by flow cytometry. (G) Mitochondrial activity and (H) mass were determined by the TMRE and MitoTracker Green probes, respectively, and flow cytometry. (I) Quantification of glucose uptake via flow cytometry using a fluorescent glucose analog. (J) Flow cytometric detection of newly synthesized proteins using Click-iT Plus OPP Alexa Fluor 488 Protein Synthesis Assay and flow cytometry. (K) Cytosolic calcium iron (Ca^2+^) levels in RPMs of IB and ID mice were measured using Cal-520 fluorescent probe with flow cytometry. (L) Cytosolic Ca^2+^ levels in RPMs of IB and ID mice were measured with flow cytometry post-PR73 administration using Cal-520 fluorescent probe with flow cytometry. (M) Cytosolic Ca^2+^ levels in iRPMs cultured with serum from IB or ID mice and serum from ID mice treated with the hepcidin agonist PR73 (2 µg/mL, 4 h) were determined using Cal-520 fluorescent probe with flow cytometry. (N) Flow cytometric measurement of cytosolic Ca^2+^ levels in mock and FPN-iRPMs with and without sRBC exposure using Cal-520 fluorescent probe with flow cytometry. (O) Normalized EP capacity of PKH26-sRBCs by mock and FPN-iRPMs treated with an intracellular (BAPTA-AM; 10 µM, 1 h) and extracellular (EGTA; 3 mM, 30 min) calcium chelator. (P) The EP capacity in iRPMs cultured with serum from IB and ID mice, and treated with extracellular (EGTA; 3 mM, 30 min) calcium chelator, was determined using flow cytometry by measuring the percentage of cells that phagocytosed PKH26-labeled sRBCs. (Q) Flow cytometric assessment of EP capacity in mock-transduced iRPMs and iRPMs overexpressing wild-type FPN and FPN mutant D39A was quantified as the percentage of cells engulfing PKH26-labeled RBCs. (R) Flow cytometric measurement of cytosolic Ca^2+^ levels in mock, FPN, and D39A-iRPMs using Cal-520 fluorescent probe with flow cytometry. Each dot represents one mouse or an independent cell-based experiment. Data are represented as mean ± SEM. Welch’s unpaired t-test determined statistical significance in A-C and K. Two-way ANOVA with Tukey’s Multiple Comparison tests was used in D-J, N, and N-P. A one-way ANOVA test with Dunnett’s or Tukey’s Multiple Comparison tests was used in M and Q-R. ns p>0.05, *p<0.05, **p<0.01, ***p<0.001 and ****p<0.0001.

We next investigated in more detail the potential metabolic consequences of increased FPN expression. FPN overexpression led to a significant increase in both mitochondrial activity and mass compared to mock-transduced iRPMs in a manner that was likewise blocked by vamifeport (Figure 4G-H). This response was reflected by a vamifeport-controlled increase in glucose uptake and protein synthesis rate in FPN-overexpressing iRPMs (Figure 4I-J), suggesting a coordinated metabolic adaptation. Importantly, these responses were independent of the presence of sRBCs in the culture media and intracellular iRPM LIP levels (Figure S4A-D), indicating a regulatory effect of FPN on mitochondrial functions that may reach beyond its role as a regulator of intracellular iron status.

We next set out to elucidate the signaling mechanisms underlying the functional rewiring of RPMs in response to increased FPN expression during systemic iron deficiency. Our previous work demonstrated that age-related iron overload, likely due to low FPN expression, suppressed RPM’s RBC clearance capacity^27^. This phenomenon was recapitulated in iron-loaded *in vitro* iRPMs and was reflected by reduced intracellular Ca^2+^ levels^27^, a key regulatory signal for macrophage phagocytosis^32,59^. Extending this observation, we found a significant increase in intracellular Ca^2+^ levels within RPMs of ID mice compared to IB mice (Figure 4K), mirroring the enhanced RBC clearance capacity. Notably, PR73 administration reversed the elevated intracellular Ca^2+^ levels specifically in RPMs of ID mice, with no effect observed in RPMs of IB mice (Figure 4L). In parallel, iRPMs exposed to ID serum also displayed increased Ca^2+^ levels, a response likewise suppressed by PR73 treatment (Figure 4M). This demonstrates that intracellular Ca^2+^ signaling, like phagocytic and mitochondrial activities, is uniquely reprogrammed in RPMs of iron-deficient mice in a manner dependent on low hepcidin/high FPN. Consistently, FPN-iRPMs exhibited increased intracellular Ca^2+^ levels compared to mock-iRPMs, independent of the presence of stressed RBCs during culture and cellular LIP levels, further implying the role of Ca^2+^ signaling in RPM functional rewiring driven by high FPN (Figure 4N). To further examine whether enhanced Ca^2+^ signaling may support increased EP in iRPMs, we employed both intracellular (BAPTA-AM) and/or extracellular (EGTA) Ca^2+^ chelators in the FPN-iRPM model and iRPMs exposed to ID and IB serum (Figure 4O-P). Both chelation strategies not only reduced the overall EP capacity of iRPMs but also effectively suppressed the enhancement of RBC clearance in FPN-high conditions. Recent findings implied that FPN may have the capacity for Ca^2+^ import^60^. To test whether FPN-mediated Ca²^+^ influx directly drives their functional rewiring, we employed overexpression of the FPN variant harboring a specific aspartate-to-alanine mutation (D39A), expected to disrupt the unidirectional movement of Ca^2+^ into the cell while preserving Fe²⁺ transport function^54^. Interestingly, we found no significant differences in the enhancement of EP or intracellular Ca^2+^ levels between iRPMs overexpressing mutant D39A and wild-type FPN (Figure 4Q-R). In sum, our findings suggested that FPN-mediated activation of Ca^2+^ signaling, likely through an indirect mechanism, contributes to enhanced phagocytic capacity of RPMs.

### SYK kinase mediates the FPN-dependent adaptation of RPMs to ID conditions

Expanding on prior knowledge, we next sought to identify the genetic mediators of Ca^2+^-dependent signaling underlying the functional rewiring of RPMs in ID mice. First, we examined the involvement of PIEZO1, a mechanoreceptor identified to modulate Ca^2+^ signaling and EP in RPMs^32^. Indeed, under basal conditions, treatment of iRPMs *in vitro* with the PIEZO1 activator YODA1 and the inhibitor GsMTx1 mildly affected EP (Figure S5.1A-B). However, GsMTx1 did not significantly affect the enhanced EP capacity observed in iRPMs exposed to ID serum (Figure S5.1C). These findings excluded a major role for PIEZO1 in mediating the functional acceleration of RPMs in response to iron deficiency.

To further dissect the Ca^2+^-dependent signaling pathways responsible for the enhanced EP observed in RPMs during iron deficiency, we performed a targeted pharmacological screen. Inhibitors of CaMKK (STO-609), calmodulin-mediated signaling (W7), P2 receptor-mediated Ca^2+^ signaling (Apyrase)^59^, and macropinocytosis (EIPA) had no discernible impact on EP in either mock or FPN-iRPMs (Figure 5A). Likewise, despite its well-established role in Ca^2+^-dependent signaling, inhibition of protein kinase C (PKC) with Bisindolylmaleimide I did not reduce EP in either mock-or FPN-iRPMs (Figure 5A). However, the inhibition of the spleen tyrosine kinase (SYK), a downstream effector of PKC-dependent signaling, with piceatannol completely abolished the increased EP in FPN-iRPMs *versus* mock-transduced iRPMs (Figure 5A). To mitigate the potential off-target effects of piceatannol, we next employed the highly selective SYK inhibitor entospletinib (ENTO; also known as GS-9973) in mock- and FPN-iRPM cultures. Consistently, ENTO effectively suppressed FPN-driven EP enhancement in FPN-iRPMs and iRPMs cultured in ID versus IB serum (Figure 5B-C). Furthermore, we observed that ENTO prevented an increase in Ca²⁺ levels upon FPN overexpression (Figure 5D). Interestingly, supplementation with ENTO was sufficient to reduce both the mitochondrial activity and mass of iRPMs overexpressing FPN to the control levels of mock-transduced cells (Figure 5E-F).

**Figure 5.**
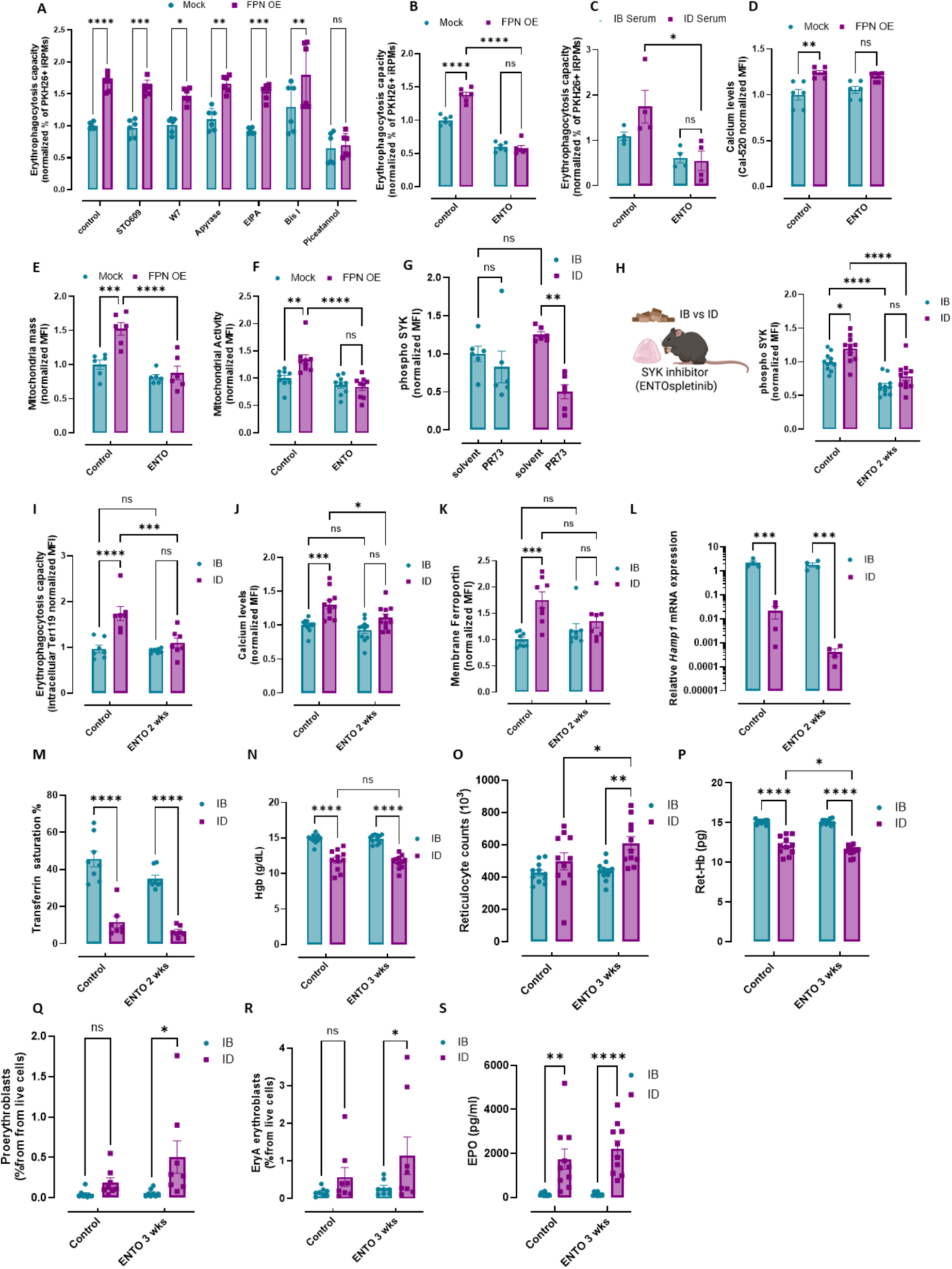
SYK kinase mediates the FPN-dependent adaptation of RPMs to ID conditions. (A) Normalized EP capacity of mock and FPN-iRPMs, measured by uptake of PKH26-labeled sRBCs with flow cytometry. iRPMs were cultured with sRBC for 4 days before analysis and treated with inhibitors targeting CaMKK (STO-609; 2.5 µM, 16 h), ATP-triggered purinergic receptor-mediated Ca²⁺ signaling (Apyrase; 5U/ml, 30 min), calmodulin (W7; 20 µM, 30 min), macropinocytosis (EIPA; 25 µM, 30 min), protein kinase C (Bisindolylmaleimide I, BIS I; 10 µM, 1 h), and spleen tyrosine kinase (SYK; Piceatannol; 100 µM, 30 min). (B) Normalized EP capacity of mock and FPN-iRPMs upon treatment with the SYK inhibitor Entospletinib (ENTO) measured by uptake of PKH26-labeled sRBCs. (C) Normalized EP capacity of iRPMs cultured with serum from IB or ID mice upon treatment with ENTO was measured by uptake of PKH26-labeled sRBCs with flow cytometry. (D) Flow cytometric measurement of cytosolic Ca^2+^ levels in mock and FPN-iRPMs upon treatment with ENTO using Cal-520 fluorescent probe. (E) Mitochondrial mass was determined in mock and FPN-iRPMs upon ENTO treatment using MitoTracker Green and flow cytometry. (F) Mitochondrial activity was measured in mock and FPN-iRPMs with and without treatment with ENTO using the TMRE probe and flow cytometry. (G) Flow cytometric analysis of phosphorylated SYK (phospho-SYK) in RPMs isolated from IB and ID mice, 4 h after intraperitoneal administration of the hepcidin agonist PR73 (50 nmol/mouse). (H). IB and ID mice were supplemented with the orally bioavailable SYK inhibitor ENTO via daily jelly feeding for 2 or 3 weeks before analysis (5 mg/kg/day). Phospho-SYK levels in RPMs were assessed by flow cytometry in IB and ID mice supplemented with ENTO. (I) EP capacity of RPMs from IB and ID mice supplemented with ENTO was determined by intracellular staining for the erythrocyte marker TER119, followed by flow cytometry. (J) Cytosolic Ca²⁺ levels in RPMs from IB and ID mice supplemented with ENTO were measured by flow cytometry using the Cal-520 fluorescent probe. (K) Surface expression of FPN on RPMs was assessed by flow cytometry in IB and ID mice supplemented with ENTO. (L) Relative mRNA expression of the Hamp gene in RPMs from IB and ID mice supplemented with ENTO. (M) Plasma transferrin saturation levels were measured in IB and ID mice supplemented with ENTO. (N) Blood hemoglobin (Hgb) concentration was determined in IB and ID mice supplemented with ENTO. (O) Reticulocyte counts in IB and ID mice supplemented with ENTO. (P) Reticulocyte hemoglobin content in IB and ID mice supplemented with ENTO. (Q and R) Proerythroblast (Q) and EryA erythroblasts (R) representations in the spleens of IB and ID mice supplemented with ENTO. (S) Plasma EPO levels were measured in IB and ID mice supplemented with ENTO. Each dot represents either an independent cell-based experiment or one mouse. Data are represented as mean ± SEM. Two-way ANOVA with Tukey’s Multiple Comparison tests was used to determine statistical significance. ns p>0.05, *p<0.05, **p<0.01, ***p<0.001 and ****p<0.0001.

Next, we sought to investigate the role of SYK in RPM iron recycling *in vivo.* First, we found that a 4 h treatment of ID mice with PR73 markedly reduced levels of the active form of SYK, phospho-SYK (Y525/526), in RPMs, whereas no significant effect of the hepcidin agonist was observed in RPMs from IB mice (Figure 5G), similarly to all previous ID-specific suppressive effects of PR73 on RPM rewiring phenotypes. We further assessed the impact of the SYK inhibition with ENTO in both IB and ID mice. Oral daily ENTO administration for the last 2 weeks of the iron-modified diet (5 mg/kg/day; with flavored jellies) effectively reduced phospho-SYK (Y525/526) levels in RPMs from both IB and ID mice compared to vehicle controls (Figure 5H). Strikingly, ENTO administration led to a marked reduction in the EP capacity of RPMs, specifically in ID mice, while EP in IB mice remained unaffected by ENTO treatment (Figure 5I). Furthermore, ENTO treatment partially reversed the increase in Ca²⁺ levels in RPMs observed between ID and IB mice (Figure 5J). Interestingly, ENTO also suppressed FPN induction associated with iron deficiency (Figure 5K), suggesting that EP capacity and FPN expression may be bidirectionally linked. The modulation of FPN appeared independent of liver hepcidin mRNA expression, with no hepcidin induction observed by ENTO supplementation (Figure 5L). Instead, hepcidin expression in ENTO-treated ID mice dropped approximately 50-fold compared to placebo-ID mice. This may indicate that ENTO treatment promotes hepcidin-suppressing signals, such as increased hypoxia or erythropoietic drive, but the exact mechanism by which SYK regulates hepcidin remains to be determined. On systemic levels, a 2-week ENTO treatment did not affect liver iron content within IB or ID groups (Figure S5.2A) or circulating hematological parameters (Figure S5.2B-C), and it only showed some tendency for the more pronounced decrease in the transferrin saturation levels (Figure 5M). Therefore, we extended the period of ENTO administration for 3 weeks and evaluated the systemic blood parameters. While we observed no significant difference in hemoglobin or MCH between iron-restricted mice supplemented with ENTO *versus* placebo (Figure 5N, S5.2D), we noticed an increase in reticulocyte counts and reduction of reticulocyte hemoglobin content (Figure 5O-P), indicating the onset of systemic hematological effects of ENTO treatment. Consistently, whereas erythroid progenitors’ expansion was not affected by ID in the bone marrow, we detected a more pronounced expansion of immature EPO-responsive splenic progenitors, proerythroblasts, and EryA erythroblasts, in ENTO-treated *versus* control mice (Figure 5Q-R). Consistently, plasma EPO levels were elevated in a more pronounced manner between ID and IB mice following ENTO supplementation compared to placebo-exposed mice (Figure 5S). Interestingly, the saturation of transferrin in the circulation was not significantly different between ID mice treated with ENTO and placebo (Figure S5.2E). This suggests that a more pronounced compensatory extramedullary erythropoietic response is initiated by ENTO, likely due to less effective direct iron delivery from iron-recycling RPMs. Taken together, SYK activity emerged as critical for the enhanced RBC-clearance capacity of RPMs in ID mice, with SYK inhibition triggering compensatory erythropoietic responses that are consistent with a possible reduction in iron delivery from RPMs.

### Functional adaptation of RPMs to iron deficiency involves enhanced BCAA catabolism and β-oxidation pathways

Efferocytosis is a continuous, energy-intensive process that demands extensive metabolic resources and reprogramming^5^. Enhanced glucose uptake and potentiated OXPHOS of ID *versus* IB RPMs, processes regulated in part by FPN (Figure 3N and P), have been previously reported to support efferocytosis^10,11^. However, we questioned whether the metabolic reprogramming observed in RPMs of ID mice reflects a unique adaptation to the challenge of processing hemoglobin-rich RBCs or if it shares features with other macrophage efferocytic programs. To address this, we revisited our transcriptomic and proteomic datasets. Morioka *et al*. identified nineteen solute carrier (SLC) transporter genes that are transcriptionally upregulated during apoptotic cell clearance, establishing a canonical efferocytosis signature^10^. Strikingly, RPMs in ID mice diverge from this model: none of these nineteen SLC transporters were upregulated at the mRNA level. Similarly, we observed no clear signature of hypoxia-driven programs, pentose phosphate pathway activation, or increased lactate production^9,61^. Notably, RPMs also failed to display detectable protein levels of key enzymes associated with amino acid signaling that support efferocytosis, including ARG1 and ODC1 (crucial for arginine-to-putrescine conversion), or IDO1 (the tryptophan-catabolizing enzyme), and did not express SLC36A4, the principal tryptophan transporter^16,18^. While GLS1, the enzyme responsible for non-canonical glutaminolysis^19^, was detected through proteomic analysis, its levels were unaltered under iron-deficient conditions. Conversely, the enriched category of metabolic proteins differentially regulated in RPMs of ID mice revealed consistent alterations of lipid metabolism, resembling other efferocytic macrophages^11^. This includes elevated enrichment expression of β-oxidation enzymes, most notably the rate-limiting CPT2, induction of lipases, and concurrent downregulation of DGAT1 (diacylglycerol O-acyltransferase 1), which mediates triacylglyceride synthesis. Interestingly, we also observed significant enrichment of enzymes involved in branched-chain amino acid (BCAA) breakdown. In parallel, two upregulated proteomic hits, SLC7A7 and SLC7A8, represent antiporters that transport cationic and large neutral amino acids, including BCAAs^62^.

To validate these metabolic adaptations in RPMs of ID mice and assess whether they are specific to this subset, we profiled selected proteomic hits by flow cytometry across tissue-resident macrophage populations. We found that BCKDHB, the rate-limiting enzyme of BCAA catabolism, the BCAA exporter SLC7A7, β-oxidation enzymes CPT2 and ACADSB, the TCA enzyme citrate synthase, and the lipase ABHD12B were all significantly elevated in ID RPMs compared to iron-balanced controls, while remaining unchanged or reduced in KCs and peritoneal macrophages (Figure 6A). Consistently, both DGAT1 and lipid droplet levels were specifically downregulated in RPMs of ID mice, suggesting a unique skewing toward energy-consuming pathways under iron deficiency (Figure 6A). Next, we focused on BCAA metabolism, a pathway not previously linked to enhanced efferocytosis. Baseline BCAA levels were exceptionally high in RPMs compared to other splenic cell types, though still lower than in KCs and similar to peritoneal macrophages (Figure 6B-C). BCAA content increased significantly in RPMs treated *ex vivo* with ERG240, a pharmacologic blocker of BCAA catabolism at the stage of the first reaction catalyzed by the BCAT enzyme (Figure 6C). Prolonged exposure of iRPMs to stressed RBCs, but not RBC ghosts, induced strong BCKDHB expression, more robust than in standard BMDMs, and appeared to prevent BCAA accumulation in RBC-exposed cells (Figure 6D-E). Notably, BCAA content remained unchanged in RPMs from ID mice (Figure 6F), despite elevated EP, suggesting that BCAA metabolism is tightly regulated. This implied that ID conditions impose a new metabolic equilibrium necessary to maintain enhanced iron recycling by RPMs. Further exploring this concept, BCKDHB levels were elevated in iRPMs overexpressing FPN, a response that, like EP, depended on both FPN and SYK kinase activity (Figure 6G). The SYK dependency of BCKDHB was also evident *in vivo* in RPMs derived from ID *versus* IB mice (Figure 6H). Strikingly, inhibition of BCAT enzymatic activity using ERG240 abolished the EP boost caused by FPN overexpression (Figure 6I), indicating that enhanced BCAA catabolism is essential for reprogramming RPM phagocytic function. Similarly, exposure to leucine increased EP capacity in iRPMs in a manner diminished by ERG240 pre-treatment (Figure 6J). Interestingly, BCAT inhibition also reversed BCKDH induction and reduced mitochondrial mass and activity, specifically in FPN-overexpressing iRPMs (Figure 6K-M). We confirmed the reversal of the enhanced phagocytic and metabolic function of ID-like RPMs by BCAT suppression using another inhibitor, BAY069 (Figure S6.1A-C). We next looked at the requirement of β-oxidation for the RPM ID-like adaptation. Like BCKDHB, CPT2 upregulation was dependent on both FPN and SYK activity (Figure S6.2A). This SYK dependence was further validated *in vivo*, where ENTO reversed CPT2 induction in RPMs from ID mice (Figure S6.2B). Finally, inhibition of CPT2 activity with trimetazidine (TMZ) abolished the enhanced EP observed with FPN overexpression, demonstrating that increased β-oxidation is a key component of the metabolic programming driving augmented phagocytic function in RPMs (Figure S6.2C). Taken together, both BCAA catabolism and β-oxidation emerged in our study as FPN/SYK-controlled pathways that maintain enhanced phagocytosis and metabolic activities of RPMs in iron deficiency.

**Figure 6.**
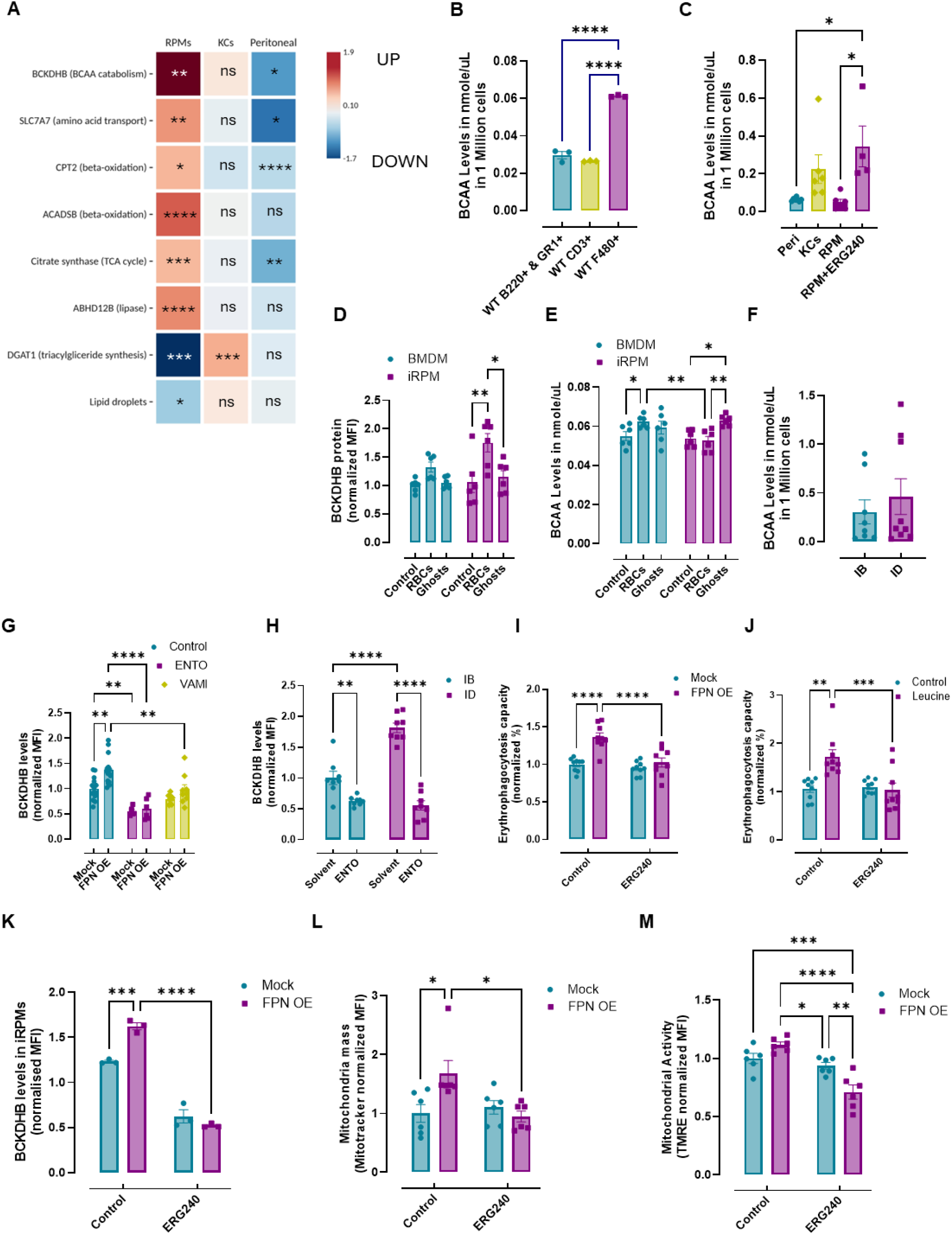
Functional adaptation of RPMs to iron deficiency involves enhanced BCAA catabolism. (A) Heatmap showing levels of BCKDHB, SLC7A7, CPT2, ACADSB, citrate synthase, ABHD12B, DGAT1, and lipid droplet levels across RPMs, KCs, and peritoneal macrophages measured by flow cytometry. Data represent log2-transformed fold changes in cells of ID mice vs IB mice, with significant changes marked by asterisks. (B) Quantification of BCAA levels in different splenic immune cell populations - F4/80⁺ RPMs, CD3⁺ T cells, and other splenocytes, measured using a colorimetric BCAA assay. (C) Quantification of BCAA levels of *ex vivo* magnetically-sorted peritoneal macrophages, KCs, and RPMs (untreated and treated with ERG240). (D) Intracellular BCKDHB levels and (E) BCAA levels in iRPMs and BMDMs treated with stressed RBCs and RBC ghosts compared to untreated control. (F) BCAA levels in magnetically-isolated RPMs from IB and ID mice (G). Intracellular BCKDHB levels in mock and FPN overexpressing (OE) iRPMs upon treatment with entospletinib (ENTO) (10 µM, 1 h) and vamifeport (VAMI) (20 μM, 15 h), determined using flow cytometry (H). Intracellular BCKDHB levels in RPMs from IB and ID mice supplemented with ENTO via jelly feeding (3 weeks) were determined using flow cytometry. (I) Normalized EP capacity towards PKH26-sRBCs by mock and FPN overexpressing iRPMs upon ERG240 treatment (10 µM, 15 h). (J) Normalized EP capacity towards PKH26-sRBCs by iRPMs upon leucine exposure. (K) Intracellular BCKDHB levels in mock and FPN OE iRPMs upon treatment with ERG240. (L) Mitochondrial mass was determined in mock and FPN OE iRPMs upon treatment with ERG240 using MitoTracker Green and flow cytometry. (M) Mitochondrial activity was determined in mock and FPN OE iRPMs upon treatment with ERG240 using the TMRE probe and flow cytometry. Each dot represents one mouse or an independent cell-based experiment. Data are represented as mean ± SEM. Two-way ANOVA with Tukey’s Multiple Comparison tests was used in D, E, and G-M, while one-way ANOVA test with Dunnett’s or Tukey’s Multiple Comparison tests was used in B and C, A Welch’s unpaired t-test determined statistical significance in A and F. ns p>0.05, *p<0.05, **p<0.01, ***p<0.001 and ****p<0.0001.

### *In vivo* inhibition of BCAA catabolism alters erythrophagocytic capacity and metabolic remodeling of RPMs in ID mice

We next sought to dissect the functional relevance of BCAA metabolism in regulating erythrophagocytic capacity and mitochondrial remodeling *in vivo*. First, we found that enhanced OXPHOS of freshly isolated RPMs from ID mice, as measured using a Seahorse analyzer, was abrogated by *ex vivo* treatment with ERG240 (Figure 7A), indicating its critical dependence on BCAA catabolism. Next, we administered ERG240, a bioavailable BCAT inhibitor, to ID mice via daily jelly feeding for three weeks. Consistent with our *in vitro* observations, ERG240 treatment significantly suppressed the enhanced EP activity of RPMs in ID mice (Figure 7B). In parallel, ERG240 administration prevented the increase of mitochondrial mass and the upregulation of BCKDHB and citrate synthase protein levels, a proxy for TCA cycle capacity (Figure 7C-E). Interestingly, despite this reduction in mitochondrial mass and citrate synthase, mitochondrial membrane potential was not suppressed and remained comparable to untreated ID controls (Figure 7F), suggesting a partial uncoupling between mitochondrial polarization state and respiratory functions under conditions of BCAT inhibition *in vivo*. Notably, FPN protein abundance remained unchanged in ERG240-treated mice subjected to ID feeding (Figure 7G), indicating that, as in ENTO-supplemented animals, reduced RPM erythrophagocytic activity is linked to diminished exposure of FPN at the plasma membrane. At the systemic level, ERG240 treatment very modestly affected the degree of anemia, as evidenced by a slightly more pronounced reduction in hematocrit, while hemoglobin levels, MCV, and MCH values remained unchanged in ID mice receiving the drug relative to untreated ID controls (Figure 7H and S7A). However, circulating reticulocyte counts were significantly elevated specifically in ERG240-treated animals (Figure 7I), indicating a heightened stress erythropoietic response aimed at restoring circulating red cell mass, likely reflecting enhanced sensing of iron deficiency. In line with this, ERG240-treated mice displayed a significant expansion of splenic erythroid progenitors, with both proerythroblasts and EryB erythroblasts (Figure S7D) showing elevated representation relative to untreated ID mice. Consistently, plasma EPO levels were significantly elevated in ERG240-treated ID mice (Figure 7J), aligning with the observed expansion of splenic erythroid progenitors. Interestingly, not only were *Epo* mRNA levels elevated in the kidney of ID mice upon ERG240 supplementation as compared to placebo controls, but also *Tfrc* transcription was increased (Fig. 7K and L). Similarly, cardiac *Tfrc* appeared as more significantly induced in ID mice upon ERG240 treatment compared with untreated ID controls, reflecting a slightly more pronounced drop in the heart non-heme iron content (Fig. S7E-F). However, liver *Tfrc* induction and a decrease in iron levels upon ID diet were not modified by ERG240 supplementation (Fig. S7G-H), and transferrin saturation remained comparably reduced in ERG240- and placebo groups (Fig. S7I). These observations may suggest sensing of more pronounced hypoxia by the kidney, and of tissue iron deficiency in some, but not all, organs.

**Figure 7.**
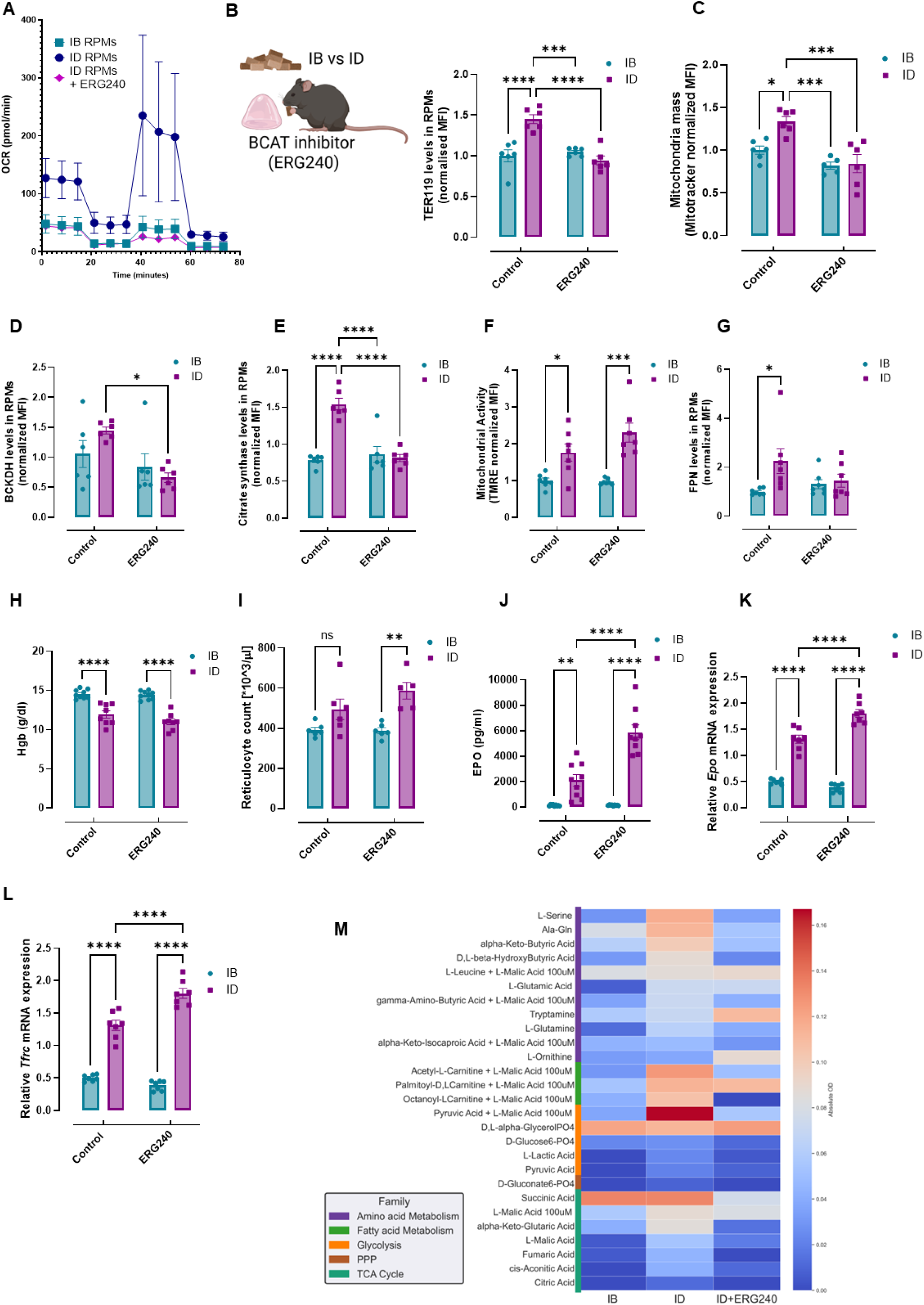
*In vivo* inhibition of BCAA catabolism alters erythrophagocytic capacity and metabolic remodeling of RPMs in ID mice. (A) Seahorse analysis of *ex vivo* magnetically-isolated RPMs from IB and ID mice, including 4 h ERG240 preincubation for RPMs from ID mice. (B) IB and ID mice were supplemented with the orally bioavailable BCAT inhibitor ERG240 via daily jelly feeding for 3 weeks before analysis (5 mg/kg/day). EP capacity of RPMs from IB and ID mice supplemented with ERG240, determined by intracellular staining for the erythrocyte marker TER119, followed by flow cytometry. (C) Mitochondrial mass of RPMs from IB and ID mice supplemented with ERG240 was determined using MitoTracker Green, followed by flow cytometry. (D) Intracellular BCKDHB levels of RPMs from IB and ID mice supplemented with ERG240 (E). Intracellular citrate synthase levels of RPMs from IB and ID mice supplemented with ERG240 (F). Mitochondrial activity of RPMs from IB and ID mice supplemented with ERG240 was determined using the TMRE probe, followed by flow cytometry. (G) FPN levels of RPMs from IB and ID mice supplemented with ERG240 were determined by flow cytometry. (H) Blood hemoglobin (Hgb) and reticulocyte count (I) were determined in IB and ID mice supplemented with ERG240. (J) Plasma EPO levels were measured in IB and ID mice supplemented with ERG240. (K and L) Relative mRNA expression of the (K). Relative mRNA expression of *Epo* and (L) *Tfrc* genes in the kidneys of IB and ID mice supplemented with ERG240 (M). Absolute mitochondrial substrate utilization profiles of RPMs from IB, ID, and ID mice supplemented with ERG240, measured using the MitoPlate S1 assay. Blank-corrected OD values for all metabolites with detectable activity in ID RPMs (n = 27) are shown and grouped by metabolic pathway. Each dot represents one mouse or an independent cell-based experiment. Data are represented as mean ± SEM. Two-way ANOVA with Tukey’s Multiple Comparison tests was used in B-L. ns p>0.05, *p<0.05, **p<0.01, ***p<0.001 and ****p<0.0001.

Finally, to assess how systemic iron deficiency alters mitochondrial fuel use *in vivo*, and how these changes depend on BCAT activity, we performed MitoPlate S1 substrate profiling on freshly isolated RPMs. Relative to IB controls, RPMs from ID mice exhibited broadly increased utilization of glycolytic, amino-acid–derived, fatty-acid/carnitine-linked, and TCA cycle substrates, consistent with their heightened OXPHOS phenotype (Figure 7M). BCAT inhibition with ERG240 partially attenuated this shift, most notably reducing oxidation of several TCA cycle and fatty-acid–linked fuels, in line with the decrease in global mitochondrial respiration measured by Seahorse. In contrast, several amino-acid substrates, such as leucine+L-malic acid, tryptamine, and ornithine, remained more highly utilized than in RPMs of IB mice, indicating that while BCAT-dependent BCAA metabolism contributes to the enhanced respiratory capacity of RPMs in ID mice, additional metabolic pathways support the full extent of their increased mitochondrial activity.

Collectively, these data demonstrate that BCAA catabolism is required for sustaining increased EP functions, with implications for the body’s adaptation to dietary iron restriction.

## Discussion

Iron deficiency is a complex physiological state with multifaceted implications. The precise quantitative thresholds that trigger cellular or organismal functional consequences, and the cell-type-specific molecular consequences, including the identification of the most vulnerable cell types and the sequence of cellular dysfunction triggered by iron deficiency, remain poorly understood. Beyond its well-known erythropoietic effects leading to anemia, iron deficits impair neuronal and muscle functions^33,63,64^, suppress immune cell activity and proliferation^65–68^, and dysregulate macrophage responses^49,50,69,70^. However, the seemingly contradictory findings regarding the effects of iron depletion on macrophage inflammatory cytokine production *in vitro* versus *in vivo*^49,50,69,70^ highlight the critical need for a more precise, cell-type-specific, and context-dependent understanding of how low iron status affects macrophage functions. Specifically, how RPMs, specialized iron-recycling cells central to the maintenance of systemic iron homeostasis, adapt to the metabolic stress of systemic iron deficiency remains a key open question. Here, we address this gap by defining a signaling-metabolic circuit through which systemic iron deficiency reprograms RPMs to enhance erythrophagocytosis and sustain iron recycling (Fig. 8).

**Figure 8.**
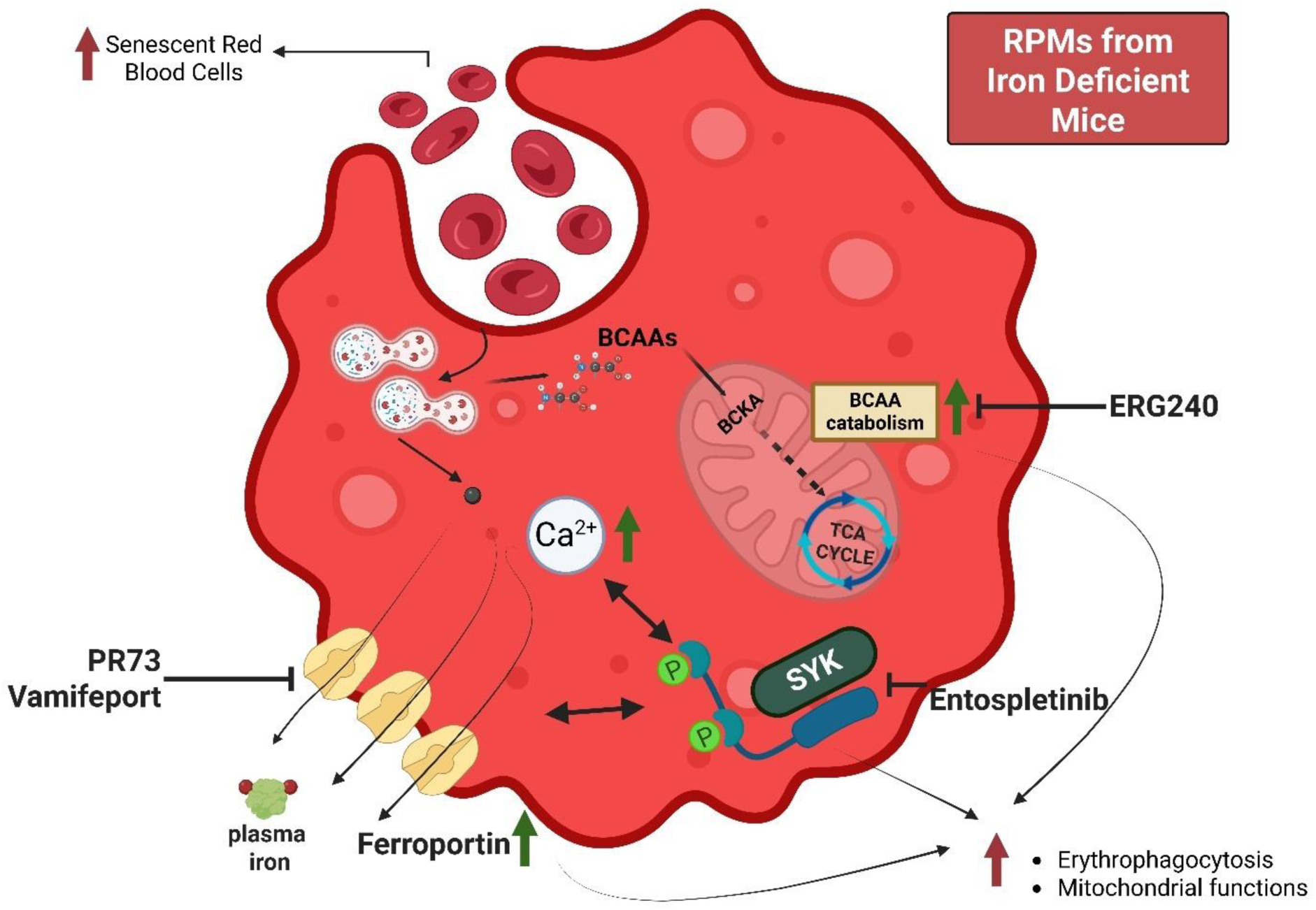
Iron deficiency induces metabolic rewiring of red pulp macrophages to enhance erythrophagocytosis. Model illustrating how systemic iron deficiency reprograms RPMs to sustain iron recycling. Increased FPN-mediated iron export activates SYK-dependent signaling, promoting mitochondrial metabolic adaptation, including BCAA catabolism, which supports enhanced erythrophagocytosis of senescent RBCs. Pharmacological inhibition of FPN, SYK, or BCAA catabolism disrupts this adaptive program.

RPMs from ID mice displayed classical markers of cellular iron depletion (increased TFR1, decreased ferritin), reflecting activation of the iron-responsive element/iron regulatory protein (IRE/IRP) system, a post-transcriptional regulatory mechanism that normally increases iron uptake and reduces iron storage and FPN-mediated export under iron-limiting conditions^20,25^. However, we observed increased FPN expression in RPMs of ID mice, despite proteomic analysis showing elevated IRP2 levels. This implies that suppressed hepcidin levels in iron deficiency, which relieve the dominant negative regulation of FPN, override any potential IRP2-mediated repression. Although RPM iron stores were low in ID mice, increased LIP and oxidative stress were observed relative to IB controls, consistent with enhanced EP activity and a likely increase in cellular iron flux. Intriguingly, this observation suggests that other iron-related signals beyond LIP contribute to IRP2 stabilization in RPMs. The elevated FPN expression facilitates the export of iron derived from engulfed RBCs, likely supporting the accelerated stress erythropoiesis in the spleen and the iron needs of other cell types during iron deficiency. These results reveal an adaptive mechanism in RPMs that prioritizes systemic iron delivery while promoting more efficient clearance of senescent RBCs in the spleen in response to dietary iron deficiency. Importantly, by demonstrating that the RBC clearance capacity of RPMs in ID mice depends on functional FPN, our findings highlight a tight coordination between iron export and its delivery through erythrophagocytosis, representing, to the best of our knowledge, a previously unrecognized concept. The relatively mild systemic effects observed upon disruption of these adaptive mechanisms may reflect the short-term nature of their experimental inhibition in RPMs. Nevertheless, our findings underscore the need to determine how RPM rewiring and increased EP during iron deficiency reshape tissue iron distribution, and how these metabolic and functional adaptations ultimately influence organismal physiology, including immune competence, neuronal function, and muscle performance.

The established cellular response to iron deficiency, observed in iron-deprived macrophages *in vitro* and in TFR1-null cardiomyocytes, involves a metabolic shift toward glycolysis accompanied by mitochondrial dysfunction and impaired electron transport chain activity^49,71^. In contrast, RPMs from ID mice show enhanced mitochondrial function, with increased mitochondrial mass and elevated OXPHOS activity confirmed by Seahorse and proteomic analyses, accompanied by an increase in mitochondrial membrane potential. This unique, cell-type-specific metabolic adaptation, absent in Kupffer cells and peritoneal macrophages, suggests that RPMs are exceptional in adjusting their physiology to increased bioenergetic demand, imposed by boosted iron recycling capability. Whether these responses are driven by the spleen’s unique microenvironment, potentially via signals from extramedullary erythropoiesis, remains to be understood. Importantly, this exceptional metabolic state of highly enhanced mitochondrial respiration was acutely shut down by inhibition of FPN with PR73, and mirrored by suppressed glucose uptake and protein synthesis *in vivo*, pointing to a novel relationship between FPN functionality and cellular metabolism in the context of heightened efferocytic activity. These findings were further supported by the similar metabolic features induced solely by FPN overexpression in the RPM *in vitro* model.

Emerging evidence links substantial metabolic reprogramming to efferocytosis. This is hallmarked by altered amino acid utilization, particularly of arginine^16^, methionine^17^, tryptophan^18^, and glutamine^19^, lactate production^9,10^, boosted pentose phosphate pathway^14,15^, increased glucose uptake and glycolysis^9,10^, fatty acid oxidation^11^, and, critically, enhanced mitochondrial respiration^12,13^, acting in concert to sustain energetically demanding efferocytosis and promote an anti-inflammatory milieu^12,13^. While RPMs of ID mice retain some of these metabolic traits, particularly enhanced OXPHOS, our omics analyses reveal a distinct divergence from canonical efferocytic programs. For example, key transporters and enzymes involved in arginine, tryptophan, and glutamine metabolism are absent or uninduced. BCAAs represent important metabolic substrates for macrophages, fueling the TCA cycle and supporting both pro-inflammatory and anti-inflammatory polarization in a context-dependent manner^72–74^. However, the relevance of BCAA catabolism in promoting efferocytosis and controlling the functions of iron-recycling macrophages has remained largely unexplored. Our study demonstrates that RPMs of ID mice uniquely upregulate BCKDHB in a manner controlled by both FPN and SYK, identifying the new mechanistic link between the capacity of BCAA breakdown and these regulators. We further show that enhanced RBC clearance in RPMs relies on BCAA-driven bioenergetics, reflected by the dependence of their heightened OXPHOS on BCAT activity. The MitoPlate profiling further clarifies this metabolic specialization. Although RPMs from ID mice display broadly elevated mitochondrial substrate use, BCAT inhibition led to a marked divergence between fuel classes: oxidation of several key TCA cycle and fatty-acid-linked substrates was substantially reduced by ERG240, whereas utilization of select amino-acid-derived fuels, particularly leucine, malic acid, tryptamine, and ornithine, remained comparatively resilient. This pattern points to a reliance on non-canonical transamination-linked anaplerosis that sustains the α-ketoglutarate (α-KG) pool. When BCAT is suppressed, the diminished α-KG supply limits TCA cycle flux, making conventional carbon inputs more vulnerable, while specific amino acid pathways able to restore or utilize α-KG through distinct enzymatic routes can still support the elevated mitochondrial state characteristic of RPMs in ID mice. Altogether, these findings reveal either a previously unrecognized specialization of RPMs for processing BCAA-rich RBCs or suggest a more general, therapeutically exploitable metabolic dependency in efferocytic macrophages. Future work will clarify whether this pathway is restricted to RPMs or is required for efficient or increased clearance of apoptotic cells in other macrophage subtypes.

Our work further identified significantly elevated intracellular Ca^2+^ levels in RPMs of ID mice, closely dependent on functional FPN and correlating with their enhanced EP capacity. While recent studies have demonstrated that PKC activity can promote FPN exposure on the cell surface in type 2 diabetes and as a response to iron loading^75^, our findings suggest that, in the context of iron deficiency, increased FPN expression and elevated intracellular Ca²⁺ may represent parallel or intersecting pathways that together facilitate enhanced EP. It is also plausible that RPMs of ID mice, characterized by low iron stores but elevated LIP, additionally regulate FPN exposure on the surface via Ca²⁺-stimulated high PKC activity. Our data suggest no direct involvement of PKC in mediating FPN-dependent and Ca²⁺-triggered regulation of EP, at least using the cellular model of RPMs. However, RPMs of ID mice treated by ENTO not only downmodulate EP but also FPN surface levels and Ca²⁺, which may represent a secondary response, possibly mediated by PKC activity.

Our discovery that SYK drives EP in RPMs aligns with its emerging roles in diverse phagocytic processes. Bian *et al*. demonstrated that SYK activation overrides Cd47-SIRPα-mediated “don’t eat me” signals to enable inflammation-driven clearance of “self” RBC^30^. Recent studies highlight SYK as a critical mediator of microglial responses to amyloid-β (Aβ) plaques in Alzheimer’s disease, where SYK deficiency impairs Aβ phagocytosis^76,77^. Our work extends this observation by identifying iron deficiency as a physiological trigger for SYK activation in RPMs, with upstream regulation by the low hepcidin-high FPN. Furthermore, our work reveals SYK-dependent metabolic-phagocytic coupling. Whereas previous findings link high SYK activity to inhibition of mitochondrial respiration^78^, we show that mitochondrial membrane potential and mass, heightened by FPN overexpression, critically depend on SYK activity. Critically, our data imply that SYK may represent a novel molecular switch for the regulation of efferocytic capacity, a concept that requires further work.

Finally, the FPN–SYK-mediated rewiring of RPMs in iron deficiency does not resemble M1/M2-like polarization. RPMs from ID mice exhibited an ambiguous pattern of surface marker alterations, indicating no consistent pro- or anti-inflammatory skewing, and no clear regulation of immune-linked mediators in omics data. They also employed a mixed functional phenotype: elevated ROS production, driven by increased labile iron, characteristic of M1 macrophages, alongside enhanced oxidative phosphorylation, mitochondrial biogenesis, and lysosomal activity, features typically associated with M2 macrophages^79–85^. Likewise, SYK has emerged as a context-dependent regulator of macrophage polarization and function, promoting fibrotic and proinflammatory programs in diabetic kidney disease and ulcerative colitis, respecitvely and enabling immunosuppressive reprogramming of tumor-associated macrophages^86,87,88^. Together with the notion that enhanced EP in RPMs only partially resembles previously reported adaptation mechanisms to efferocytosis, our work proposes that the polarization state of RPMs in iron deficiency represents a new type of adaptation, whose resemblance in other macrophage types/environmental conditions opens an avenue for future investigations.

## Methods

### Mice

Female C57BL/6 J mice were utilized for the experiments and housed under specific pathogen-free (SPF) conditions at the Experimental Medicine Centre in Bialystok, Poland. These mice were provided with either a standard iron-balanced diet (200 mg/kg, SAFE U8958P; IB diet) or an iron-deficient diet (5.9 mg/kg, SAFE U8958P; ID diet) for 5 weeks, commencing at 4 weeks of age. 9-week-old *Tmprss6* wild-type and KO mice were provided by Dr. Delphine Meynard from INSERM, Toulouse, France. Upon arrival at the local facility at the International Institute of Molecular and Cell Biology (IIMCB), mice were acclimatized briefly before being sacrificed or subjected to further procedures, if required. Ms4a3-Cre mice were obtained from Florent Ginhoux^43^, and RosaTdT reporter mice [B6;129S6-Gt(ROSA)26Sortm9(CAG-tdTomato)Hze/J JAX#: 007905] were a kind gift of Agnieszka Kobielak. Female C57BL/6-Tg(UBC-GFP)30Scha/J (UBI-GFP/BL6) mice were kindly provided by Aneta Suwińska (Faculty of Biology, University of Warsaw, Poland). For primary cell cultures, C57BL/6 J mice were maintained in the SPF facility at the Mossakowski Medical Research Institute in Warsaw, Poland.All animal experiments and protocols were approved by the respective local ethical committees in Olsztyn and Warsaw, including the decisions: Uchwala 026, WAW2/015/2019, and WAW2/048/2020). Usage of Ms4a3Cre-RosaTdT reporter mice was approved by WAW2/126/2023. For PR73 administration, mice were intraperitoneally injected with 50 nmol of PR73 for 4 h before being sacrificed. The procedure was approved by the local ethical committee in Warsaw (WAW2/150/2019). Experiments including SYK and BCAA catabolism suppression were covered by the permissions: WAW2/068/2024 and WAW2/029/2025.

### Isolation of Single-Cell Suspensions from Murine Organs

Bone marrow cells were obtained by flushing the femurs and tibias with sterile HBSS (Gibco, 14025092) using a 25G needle. Following centrifugation at 600 x g for 10 min at 4°C, the pellet containing the isolated cells was collected for further processing. The spleen was harvested and homogenized by passage through a 70 μm cell strainer (pluriSelect, 43-50070-51). Before FACS sorting, the spleen was enzymatically digested with HBSS containing 1 mg/mL Collagenase D (Roche, 11088882001) and 50 U/mL DNase I for 30 min at 37°C to facilitate cell dissociation. The resulting cell suspension was then washed with cold HBSS and centrifuged at 600 x g for 10 min at 4°C. The harvested liver was perfused using Liver Perfusion Medium (Gibco, 17701038), followed by mincing and enzymatic digestion in Liver Digest Medium (Gibco, 17703034) at 37°C for 30 min. The digested liver tissue was then pressed through a 70 μm strainer into HBSS. The resulting cell suspension was centrifuged at 50 g for 3 min to separate the hepatocyte-enriched pellet, which was discarded. The supernatant, containing non-parenchymal cells, was further centrifuged at 700 g for 15 min at 4°C. The resulting cell pellet was resuspended in a solution of 5 mL PBS with 0.5% BSA and 5 mL of 50% Percoll (Cytiva, 17-0891-01) diluted in PBS. This suspension underwent Percoll gradient centrifugation at 700 g for 30 min at 20°C to enrich specific cell populations. The peritoneal cavity was lavaged with HBSS to collect peritoneal fluid. The retrieved fluid was then passed through a 70 μm strainer to remove debris, followed by centrifugation at 600 x g for 10 min at 4°C to pellet the peritoneal cells. Blood was collected via cardiac puncture into a heparin-coated tube. The cells were separated by centrifugation at 400 g for 10 min at 4°C following a wash with HBSS. RBCs present in a single-cell suspension were lysed using 1X RBC Lysis buffer (BioLegend, 420302) for 3 min at room temperature. However, this step was excluded when analyzing erythroid cells and RBCs. Following that, the cells were washed with 1X HBSS and centrifuged at 600g for 10 min at 4℃. The resulting pellet was processed for subsequent functional assays and antibody labeling.

### Generation of iRPMs

Mononuclear cells were aseptically harvested from the femurs and tibias of mice and subsequently cultured at a concentration of 0.5 × 10^6^/1mL in RPMI-1640 medium (Gibco, 61870036) supplemented with 10% FBS (Cytiva, SV30160.03), 1X Penicillin-Streptomycin (Gibco, 15140122), and 20 ng/mL of macrophage colony-stimulating factor (MCSF, BioLegend, 576406) in a 5% CO_2_ incubator at 37℃. On days 4 and 6 of culture, the medium was replaced with fresh complete medium supplemented with 20 µM hemin (Sigma-Aldrich, 51280). In the initial experiments, we also included 10 ng/mL interleukin-33, which was later omitted as it was not critical for mimicking RPM-like phenotypes. All experiments were performed on day 8 of culture. For treatments, hemin (Sigma-Aldrich, 51280) solution was prepared with 0.15 M NaCl containing 10% NH_4_OH. For the phagocytosis assay, iRPMs were treated for 1 h with 10 µM Entospletinib (Selleck, S7523), 15 h with 10 µM ERG240 (MedChemExpress, HY-W193545A), 15 h with 10µM BAY-069 (MedChemExpress, HY-148242), 15 h with 10 mM leucine (Sigma-Aldrich, L8000), 24 h with 100 µM deferoxamine (DFE, Sigma-Aldrich, D9533), 30 min with 10 µM BAPTA/AM (CALBIOCHEM, 196419), 1 h with 3 mM EGTA (Sigma, E8145), 16 h with 2.5 µM STO609 (Sigma-Aldrich, S1318), 30 min with 20 µM W7 (Sigma-Aldrich, 681629), 30 min with 5U/ml apyrase (Sigma-Aldrich, A6535), 30 min with 25 µM EIPA (Selleckchem, S9849), 1 h with 10 µM BisindolylmaleimideI (Sigma-Aldrich, 203290), 30 min with 100 µM Piceatannol (Sigma-Aldrich, P0453). Vamifeport was a kind gift from CSL Vifor and was purchased from MedChem Express (VIT-2763), and was used for 15 h at 20 µM concentration.

### Flow Cytometric Analysis and Cell Sorting

Cell suspensions of spleens and livers at a concentration of approximately 1 × 10^7^, along with iRPMs, were stained with LIVE/DEAD Fixable Aqua/Violet (Invitrogen, L34966/L34964) following the manufacturer’s instructions to identify dead cells. After extensive washing, Fc block was added to the cells at a 1:100 dilution in FACS buffer and incubated at 4°C for 10 min. The cells were then stained with fluorophore-conjugated antibodies at a dilution of 1:100 to 1:400, depending on the titration, in FACS buffer for 30 min at 4°C. The cells were washed thoroughly with FACS buffer and analyzed using flow cytometry. For the analysis of splenic RPM populations, cells were initially identified as CD45^+^ F4/80^high^ CD11b^dim^ TREML4^+^ after excluding lineage-positive cells (CD3^+^ T cells, B220^+^ B cells, Gr-1^+^ neutrophils/monocytes), as previously described^27^. However, since we observed that the majority of RPMs were TREML4^+^, we omitted TREML4 from the panel in further analyses. Surface antibodies against CD45 (Biolegend, 30-F11), CD45R/B220 (Biolegend, RA3-6B2), Ly-6G/Ly-6C (Gr-1) (Biolegend, RB6-8C5), CD3 (Biolegend, 17A2), F4/80 (Biolegend, BM8), CD11b (Biolegend, M1/70), and TREML4 (Biolegend, 16E5) were used. For erythroid cell analysis, TER-119 (Biolegend) and CD71 (Biolegend, RI7217) were included. Staining for proteins linked to amino acid and lipid metabolism utilized antibodies against BCKDHB (Santa Cruz Biotechnology, sc-374630), SLC7A7 (Invitrogen, PA5-113527), CPT2 (Novus Biologicals, NBP1-85471), ACADSB (Proteintech, 13122-1-AP), ABHD12B (Proteintech, 20623-1-AP), citrate synthase (Proteintech, 16131-1-AP), and DGAT1 (Novus Biologicals, NB110-41487). The analysis of liver KCs and iRPM populations involved the use of surface antibodies, including CD45 (Biolegend, 30-F11), F4/80 (Biolegend, BM8), and CD11b (Biolegend, M1/70). FPN detection was performed using a non-commercial antibody that recognizes the extracellular loop of mouse FPN (rat monoclonal, Amgen, clone 1C7) directly conjugated with Alexa488 (Alexa Fluor 488 Labeling Kit ab236553). To analyze RBCs, surface antibodies such as CD45 (Biolegend, 30-F11), TER-119 (Biolegend), and CD71 (Biolegend, RI7217) were used. Lastly, the analysis of peritoneal macrophages involved the use of surface antibodies, including CD45 (Biolegend, 30-F11), F4/80 (Biolegend, BM8), MHCII (Biolegend, M5/114.15.2), and CD11b (Biolegend, M1/70). Events were acquired using Aria II (BD Biosciences) or CytoFLEX (Beckman Coulter) and were analyzed using FlowJo or CytExpert, respectively. For RNA sequencing (RNA-seq) and ATP levels quantification, RPMs were sorted into Trizol-LS (Invitrogen, 10296010) or assay buffer, respectively, using an Aria II cell sorter (BD Biosciences) with an 85 μm nozzle.

### Functional Assays and Detection of Intracellular Fe^2+^ Iron

To assess intracellular ROS levels, CellROX Deep Red Reagent (Invitrogen, C10422) was used, and fluorescence was measured in the APC channel following the manufacturer’s protocol. The lysosomal activity was evaluated using the Lysosomal Intracellular Activity Assay Kit (Biovision, K448) and fluorescence was measured in the FITC channel as directed by the manufacturer. To measure mitochondrial activity, the tetramethylrhodamine ethyl ester (TMRE) fluorescent probe (Sigma-Aldrich, 87917) was utilized at a concentration of 400 nM for 30 min at 37 °C, and the fluorescence was measured in the PE channel. Mitochondrial mass was measured by fluorescence levels upon staining with MitoTracker Green (Invitrogen, M7514) at 100 nM for 30 min at 37 °C in the FITC channel. Intracellular calcium (Ca^2+^) levels were assessed using the Cal-520 AM probe (Abcam, ab171868) at 5 μM for 1 h at 37 °C. Briefly, surface-stained cells were incubated with the probe in HBSS, followed by immediate flow cytometric analysis in the FITC channel without further washing. Intracellular ferrous iron (Fe²⁺) was quantified using FerroOrange (Dojindo, F374) with detection in the PE channel by flow cytometry. Surface-stained cells were incubated in HBSS containing 1 μM FerroOrange for 30 min at 37 °C and immediately analyzed without further wash steps. Lipid droplet content was quantified using the Lipid Droplet Assay Kit Deep Red (Dojindo, LD06-10) following the manufacturer’s protocol, with fluorescence recorded in the APC channel. Glucose uptake was measured using the Glucose Uptake Capacity Assay Kit (Dojindo, UP03), and fluorescence was measured in the APC channel as per the manufacturer’s protocol. To perform intracellular staining, cells that had already been surface-stained were fixed using 4% PFA and permeabilized with 0.5% Triton-X in PBS. The cells were then stained with appropriate primary antibodies for 1 h at 4°C, followed by a 30-minute incubation with Alexa Fluor 488 or Alexa Fluor 647 conjugated anti-Rabbit IgG (1:1000 Thermo Fisher, A-21206). Afterward, the geometric mean fluorescence intensities (MFI) of the probes/target protein levels were measured by flow cytometry using Aria II (BD Biosciences) or CytoFLEX (Beckman Coulter). Data were then analyzed using FlowJo or CytExpert software, respectively. The MFI of the adequate fluorescence minus one (FMO) control was subtracted from the samples’ MFI to quantify the data. The data were further normalized for analysis.

### ATP levels

The levels of cellular ATP in FACS-sorted RPMs (10000 cells per sample) were measured using the ATP Assay Kit (Sigma-Aldrich, MAK473) according to the manufacturer’s instructions.

### Magnetic Sorting of RPMs

60 x 10^6^ cells were incubated for 15 min at 4℃ in PBS containing 5% Normal Rat Serum (Thermo Scientific, 10710C) and anti-CD16/32 antibody (Biolegend, 101320) in a 5 mL round-bottom tube. The cells were then labeled with anti-F4/80 (APC, Biolegend, 123116), anti-Ly-6G/Ly-6C (Gr-1) (Biotin, Biolegend, 108403), anti-CD3 (Biotin, Biolegend, 100243), anti-mouse Ly-6C (Biotin, Biolegend, 128003), and anti-B220 (Biotin, Biolegend, 103204) for 20 min at 4°C in the dark. The cells were washed with a cold PBS solution containing 2 mM EDTA and 0.5% BSA (hereafter referred to as “sorting buffer”) and centrifuged at 600 g for 10 min. The pellet was resuspended in 400 μL of sorting buffer containing 50 μL of MojoSort Streptavidin Nanobeads (Biolegend, 480016) and kept in the cold and dark for 15 min. After incubation, an additional 400 μL of sorting buffer was added to the cells, and the tube was placed on an EasyEights EasySep Magnet (STEMCELL, 18103) for 7 min. The supernatant was transferred to a fresh 5 mL tube and centrifuged at 600 g for 10 min. The pellet was resuspended in 100 μL of sorting buffer, and 10 μL of MojoSort Mouse anti-APC Nanobeads (Biolegend, 480072) were added. The suspension was gently pipetted and incubated for 15 min at 4°C in the dark. The tube was then placed on a magnet for 7 min, and the supernatant was saved for analysis. The beads with attached F4/80+ cells were washed with sorting buffer and counted under a light microscope with a Neubauer chamber. The cells were pelleted and frozen in liquid nitrogen for further analysis.

### Magnetic Sorting of RBCs, T Cells, B Cells, Granulocytes, Peritoneal Macrophages, and Kupffer Cells

RBCs, T cells, B cells, and granulocytes were magnetically isolated from spleen using the same general workflow as described for splenic RPM isolation, with modified antibody labeling and bead combinations. Single-cell suspensions were incubated with anti-CD16/32 to block Fcγ receptors on leukocytes prior to staining. Cells were stained with biotinylated antibodies against TER-119 (RBCs), B220 (B cells), and Gr-1 (granulocytes) and a PE-conjugated anti-CD3 antibody (T cells) Biotin-labeled RBCs, B cells, and granulocytes were isolated using streptavidin Nanobeads, and CD3⁺ T cells using anti-PE Nanobeads. This strategy yielded highly purified fractions of RBCs, T cells, B cells, and granulocytes for downstream analyses. Peritoneal macrophages were isolated using the same marker and bead strategy as splenic macrophages, with biotinylated antibodies against CD3, B220, and Gr-1 for lineage depletion, followed by positive selection of F4/80⁺ cells using anti-APC nanobeads.

Magnetic sorting of Kupffer cells was performed according to a two-step liver perfusion protocol with minor modifications. Briefly, mice were euthanized, and livers were perfused in situ through the inferior vena cava with Liver Perfusion Medium for 3 min, followed by perfusion with prewarmed Liver Digest Medium for 15–20 min after transection of the portal vein. Digested livers were gently dissociated in RPMI 1640, filtered through a 100-µm strainer, and subjected to two rounds of low-speed centrifugation (50 g, 3 min) to remove hepatocytes. The resulting non-parenchymal cell fraction was centrifuged at 650 g for 15 min at 4 °C, treated with red blood cell lysis buffer, and further purified using a two-layer Percoll gradient. Cells collected from the interphase between 25% and 50% Percoll were washed, Fc-blocked, and incubated with anti-F4/80 MicroBeads UltraPure. F4/80⁺ Kupffer cells were isolated using magnetized LS separation columns, washed, and prepared for downstream analyses. All cell isolations were performed at 4 °C, and purified cells were counted, pelleted, and snap-frozen in liquid nitrogen prior to intracellular BCAA quantification.

### Measurement of Cellular Iron Levels

To determine the levels of total iron in magnetically sorted RPMs, the Iron Assay Kit from Sigma-Aldrich (MAK025) was employed by following the manufacturer’s instructions. The volume of buffers and reagents used in the measurement of total iron in RPMs was adjusted accordingly. The absorbance at 593 nm was recorded using a Nanodrop ND-1000 Spectrophotometer from Thermo Fisher Scientific. Iron concentrations in RPMs (ng/μL) were calculated from the standard curve and then normalized to the number of cells in each sample.

### *In Vivo* RBC Lifespan

A sterile solution of EZ-Link Sulfo-NHS Biotin (Thermo Fisher Scientific, 21217) was prepared in PBS with a final concentration of 1 mg per 100 μL. The solution was then filtered through a 0.1 μm filter (Millipore, SLVV033RS). One day before the first blood collection, 100 μL of the sterile Biotin solution was intravenously injected into the mice. Blood samples were collected from the tail vein on days 0, 5, 12, 16, and 21, using heparinized capillaries to prevent clotting. RBCs were separated by centrifugation at 400 g for 5 min at 4°C, and each sample was resuspended in 250 μL of HBSS containing 5% normal rat serum (Thermo Fisher Scientific). The suspension was then mixed with 2 μL of fluorescently labeled anti-TER-119 and streptavidin, except for the fluorescence minus one (FMO) sample in each group, where fluorescent streptavidin was omitted. After incubation at 4℃ for 30 min, the samples were centrifuged and resuspended in HBSS. Flow cytometry was used to determine the percentage of biotinylated erythrocytes.

### Preparation of Stressed RBCs for Erythrophagocytosis Assays

The preparation and staining of stressed RBCs followed a previously outlined procedure^27,39^. To obtain the RBCs, mice were humanely euthanized, and whole blood was aseptically collected via cardiac puncture into CPDA-1 solution (Sigma-Aldrich, C4431) with a final concentration of 10%. The collected whole blood was combined, and plasma was separated by centrifugation at 400 g for 15 min at 4℃, then filtered through a 0.1 μm filter and stored at 4℃. The RBCs were then suspended in HBSS, and leukocytes were eliminated using Lymphosep (Biowest, L0560-500). Following this, the cells were subjected to a 30-minute heating period at 48℃ with continuous agitation to generate stressed RBCs. To stain the sRBCs, 1×10^10^ RBCs were resuspended in 1 ml of diluent C and mixed with 1 ml of diluent C containing 4 μM of PKH-67 (Sigma-Aldrich, MINI67) or PKH-26 (Sigma-Aldrich, MINI26). The mixture was then incubated in the dark for 5 min at 37℃, and the reaction was stopped by adding 10 mL of HBSS containing 2% FCS and 0.5% BSA. Staining was followed by two washing steps with HBSS. For *in vitro* EP assays, the cells were resuspended in RPMI-1640 and counted. For the *in vivo* approach, RBCs were resuspended to 50% hematocrit in previously collected and filtered plasma.

### *In Vitro* Erythrophagocytosis

On the 8th day post-seeding, stained, and counted sRBCs were added to iRPMs on 12-well plates in 10-fold excess for 1.5 h at 37 °C and 5% CO_2_. After incubation, the cells were extensively washed with cold HBSS to eliminate any non-engulfed sRBCs. Subsequently, the cells were detached using Accutase (BioLegend, 423201), which were then transferred to a round-bottom FACS tube, washed with HBSS, and centrifuged at 600 g for 5 min. The cells were labeled with antibodies and analyzed using flow cytometry. For time-lapse confocal imaging of the execution of RBC uptake by iRPMs, sRBCs were stained with pH-sensitive pHrodo™ Deep Red Mammalian and Bacterial Cell Labeling Kit. The imaging was performed using the Opera Phenix (PerkinElmer), an automated confocal spinning disk microscope. Imaging was conducted for 120 min, with acquisition every 1.3 min.

### *In Vivo* Erythrophagocytosis

The mice were intravenously transfused with 100 μL of RBCs resuspended in plasma to 50% hematocrit. The animals were kept in cages with ad libitum access to food and water for 1.5 h before being sacrificed for organ isolation.

### Mini-Hepcidin (PR73) Preparation

An optimized protocol was developed for preparing mini-Hepcidin (PR73, a kind gift from Elizabeta Nemeth, UCLA, USA) for intraperitoneal injection in mice. Commercially available PR73 powder may not represent the pure peptide, so we adjusted the preparation based on the estimated active component content (80%). The process began by precisely weighing the PR73 powder to account for the estimated 80% active component. This powder was then dissolved in 80% ethanol to ensure complete extraction of the peptide. The solution was thoroughly mixed using a vortex mixer to achieve a homogenous suspension. Next, the solution was carefully transferred to a sterile, low-bind vial (SL-220) to minimize potential losses due to adsorption during incubation. A brief heat treatment at 37°C for a few seconds facilitated the complete dissolution of any undissolved particles. Following the heat treatment, the vial cap was removed, and the solution was aliquoted into designated culture tubes for individual injections. A Speedvac concentrator was then used to evaporate the ethanol, resulting in a concentrated PR73 gel within each tube. To achieve a final working concentration of 0.5 nmol/μl, 2 ml of sterile, injectable-grade water was added to each tube containing the PR73 gel. The tubes were vigorously vortexed to ensure complete dissolution of the gel and homogenization of the solution. For *in vivo* experiments, 100 μl of the prepared PR73 stock solution was administered per mouse via intraperitoneal injection. A solvent control solution was prepared by following the same steps outlined above but substituting the PR73 solution with only 80% ethanol. This control solution served as a baseline for comparison with the PR73-treated mice in subsequent experiments.

### Live-Cell Imaging of Erythrophagocytosis in iRPMs

To assess erythrophagocytic capacity, iRPMs were seeded in Greiner Bio-One CELLSTAR μClear™ 96-well, Cell Culture-Treated, Flat-Bottom Microplate and cultured with IB versus ID serum. iRPMs with ID serum were treated with PR73 for 4 h. Following PR73 treatment, iRPMs were exposed to temperature-stressed, pHrodo-labeled RBCs, prepared using the pHrodo™ Deep Red Mammalian and Bacterial Cell Labeling Kit (Thermo Fisher Scientific). The stressed and labeled RBCs were added to the wells, and the plate was immediately transferred to the Opera Phenix™ High-Content Screening System (PerkinElmer), maintained at 37°C with 5% CO₂. Live-cell imaging was performed for 2 h with images captured at regular intervals to monitor the uptake of RBCs, and data were analyzed using Harmony High-Content Imaging and Analysis Software (PerkinElmer).

### BCAA levels

Intracellular branched-chain amino acid content was quantified in magnetically sorted CD3^+^ T cells, B220^+^ B cells, together with GR1^+^ granulocytes, RPMs, peritoneal macrophages, and Kupffer cells, using the Branched Chain Amino Acid Assay Kit (Sigma-Aldrich, MAK003) following the manufacturer’s instructions.

### Metabolic Flux Seahorse Assay

Metabolic flux was assessed using a Seahorse XF-96 Extracellular Flux Analyzer (Agilent Technologies) Mito Stress Test (Agilent,103010-100). Magnetically sorted RPMs (8 × 10^4^ cells/well) from IB and ID mice were plated onto poly-L-lysine-coated Seahorse XF HS Miniplates (Agilent, 103725-100). Cells were incubated in Seahorse XF RPMI medium (Agilent, 103576-100) supplemented with 2 mM L-glutamine (Agilent, 103579-100), 1 mM sodium pyruvate (Agilent, 103578-100), and 25 mM glucose (pH adjusted to 7.4; Agilent 103577-100) at 37°C in a non-CO_2_ incubator for 60 min. Real-time oxygen consumption rate (OCR) was measured under basal conditions and following sequential injections of 1 µM oligomycin, 1 µM FCCP, and a mixture of 1 µM rotenone and 1 µM antimycin A.

### Mitochondrial substrate utilization (MitoPlate) Assay

Mitochondrial substrate utilization was assessed using the MitoPlate S-1 assay (Biolog, 14105) according to the manufacturer’s protocol. Magnetically sorted RPMs from IB, ID, and ID+ERG240 mice were pooled (four mice per condition), counted, and resuspended in MitoPlate assay medium before seeding at 8 × 10⁴ cells per well onto the precoated 96-well Mitoplates, containing the dye mix MC. Plates were loaded into a Tecan Infinite 200 Pro maintained at 37°C, and kinetic OD₅₉₀ measurements were collected every 10 min throughout the assay. Absolute blank-corrected OD values from the 6-h time point were used for analysis.

### Voluntary Oral Drug Administration

Strawberry-flavored gelatin jellies were prepared under sterile conditions in 96-well plates (100 µL/well), with or without ENTO and ERG240. Gelatin (8% w/v, Sigma-Aldrich, G9391) was dissolved in warm, sterile water, and strawberry flavoring essence (1% v/v, OstroVit, 5903933916217) was added. Just before gelation, ENTO or ERG240 was added to the appropriate wells of a sterile 96-well plate to achieve the desired final concentration, while control wells received vehicle only. The gelatin solution (100 µL) was then added to each well, and the plates were refrigerated at 4°C until solid. To acclimate IB and ID mice, two drug-free jellies were provided per cage daily for 2-3 days (two mice per cage). This co-housing approach minimized stress associated with individual training. Following acclimation, mice readily consumed the jellies within 15-20 min. Drug-containing jellies were administered at approximately the same time each day for two or three weeks.

### Generation of Ferroportin Overexpressing iRPMs

We successfully cloned the wild-type *Slc40a1* gene, encoding mouse ferroportin (FPN; Uniport ID: Q9JHI9), into the HIV SFFV lentiviral vector. Lentiviral production involved transfecting the chosen lentiviral vectors (FPN or Mock) along with the psPAX2 packaging vector and the pMD2.G envelope vector into HEK293T cells. The resulting viral supernatants were harvested and titrated to determine viral concentration. Subsequently, iRPMs were transduced on Day 1 with lentiviral vectors at a multiplicity of infection (MOI) of 1. On Days 4 and 6, the culture medium was replaced with fresh complete medium supplemented with 20 µM hemin. Expression levels of surface FPN were measured in each experiment using flow cytometry. This verification step confirmed successful FPN overexpression in iRPM population, allowing for functional analysis of the consequences in subsequent experiments.

### Hepcidin Measurement with ELISA

Serum hepcidin levels were measured using the Hepcidin-Murine Compete ELISA Kit (Intrinsic LifeSciences, HMC-001) following the manufacturer’s instructions. Optical density was determined with a microplate reader at a wavelength of 450 nm.

### Erythropoietin (EPO) Measurement with ELISA

Serum EPO levels were measured using the Mouse Erythropoietin/EPO Quantikine ELISA Kit (R&D Systems, MEP00B) following the manufacturer’s instructions. Optical density was determined with a microplate reader at a wavelength of 450 nm.

### Transferrin Saturation and Tissue Iron Measurements

The levels of serum iron and unsaturated iron-binding capacity were quantified using SFBC (Biolabo, 80008) and UIBC (Biolabo, 97408) kits per the manufacturer’s instructions. Transferrin saturation was determined using the formula SFBC/(SFBC+UIBC) x 100. Tissue non-heme iron content was measured using the bathophenanthroline method, and the results were calculated based on tissue dry weight, as previously described^89^.

### RT-qPCR

RNA from cell pellets was isolated using an RNA purification kit (Eurex) according to the manufacturer’s instructions. RNA from liver, heart and kidney tissues was isolated using TRIzol™ reagent according to the manufacturer’s instructions. 400 ng of RNA was subjected to reverse transcription. In the first stage of the reaction, RNA was denatured, and random primers were attached. For this purpose, a mixture consisting of 400 ng RNA, 1 μL of random primers at a concentration of 200 ng/μL, and nuclease-free water was prepared so that the total reaction volume was 17.7 μL. The mixture was incubated at 70°C for 10 min and then cooled to 4°C. The next step was to add the reaction mixture to the tubes, which contained: 0.3 μl RevertAid H Minus reverse transcriptase, 1 μL of 10 mM dNTP solution, and 5 μL of reaction buffer. The total reaction volume was 25 μL. The reaction was carried out for 60 min at 42°C, 70°C, 10 min. The obtained cDNA was diluted 9x and then used for quantitative polymerase chain reaction (qPCR). The reaction mixture consisted of 4 μL cDNA, 7.5 μL SG qPCR Master Mix (2x), and 1.5 μL of a 5 μM solution of forward and reverse primers. The whole was supplemented with nuclease-free water to a final reaction volume of 15 μL. The qPCR reaction was carried out using the LightCycler 96 Real-Time PCR System (Roche). The reaction was pre-incubated at 95°C for 600 s, and then two-step amplification was performed 40x (95°C x 15 s, 60°C x 60 s).

### Immunohistochemistry of Spleen Tissues

The spleen was fixed in 4% PFA at 4°C for 24 h. Tissues were washed with PBS (3x30min) and incubated for 2 h in 12.5% and then in 25% sucrose solution for 48 h. Subsequently, spleen tissue fragments were embedded in the Cryomatrix medium, frozen in liquid nitrogen, and cut into 10 μm thick sections using a cryo microtome (Leica). The sections were mounted on glass slides and stored at -20°C. Before staining, the sections were warmed at room temperature for 30 min, and then their edges were surrounded using a Pap Pen Liquid Blocker. Next, the slides with sections were washed in PBS for 10 min and permeabilized in 0.1% Triton X-100 solution for 20 min. Nonspecific antibody binding was blocked by incubating the sections in 3% BSA solution for 1 h at room temperature. To visualize RPMs, the tissues were incubated with FITC-conjugated F4/80 antibody (1:100), and for FPN, with Alexa Fluor 647-conjugated FPN antibody (1:100) in 3% BSA for 2 h at room temperature in the dark. After incubation with antibodies, the tissues were washed 5 x 5 min in 0.1% Triton X-100 solution to remove unbound antibodies. To stain cell nuclei, the sections were incubated with Hoechst 33342 diluted in 0.1% Triton X-100 solution (final concentration 1 μg/ml) for 10 min at room temperature in the dark. Then, the sections were washed for 3 x 5 min in 0.1% Triton X-100 solution, incubated in PBS for 10 min, and mounted with a coverslip using ProLong™ Glass Antifade Mountant. Samples were imaged on a confocal microscope with a Plan-Apochromat 40x/1.3 NA oil immersion objective (Zeiss) LSM 800.

### Histological and Histochemical Analysis

Following fixation in 10% formalin for 24 h, spleens were stored in 70% ethanol before further preparation. The tissue was embedded in paraffin and 7 μm cross-sections were cut with a microtome (Reichert-Jung, Germany). Sections were prepared as described above. After mounting on glass slides, sections were deparaffinized, incubated with a working solution containing Perls’ Prussian Blue for 30 min, counterstained with pararosaniline solution for 2 min, and analyzed under standard light microscopy (Olympus CH2).

### Transcriptome analysis by RNA-seq

RNA-seq libraries were generated from FACS-sorted RPMs (minimum 100,000 cells per sample) using the Smart-seq2 protocol^90^, which is optimized for low-input total mRNA sequencing. RNA integrity and library quality were checked by a Bioanalyzer. Sequencing was performed on an Illumina NextSeq500 platform, producing 75-bp single-end reads at ∼50 million reads per sample. RNA-seq was carried out at the EMBL GeneCore facility (Heidelberg, Germany). Sequence reads were trimmed to remove possible adapter sequences and nucleotides with poor quality using Trimmomatic v.0.36. The trimmed reads were mapped to the Mus musculus GRCm38 reference genome available on ENSEMBL using the STAR aligner v.2.5.2b. Unique gene hit counts were calculated by using featureCounts from the Subread package v.1.5.2. Normalization and differential expression analysis were performed using DESeq2. The Wald test was used to generate p-values and log2 fold changes. Genes with an adjusted p-value < 0.05 and absolute log2 fold change > 1 were called as differentially expressed genes.

### Proteomic analysis of RPMs

#### Sample Preparation

Magnetically isolated RPMs were obtained from the spleens of IB and ID female C57BL/6J mice. The cells were lysed using RIPA buffer. Proteins were precipitated using a chloroform/methanol method, washed with methanol, and reconstituted in 100 mM HEPES (pH 8.0) containing 10 mM TCEP and 10 mM chloroacetamide.

#### LC-MS/MS Analysis

10 µg of each lysate was subjected to an in-solution tryptic digest using a modified version of the Single-Pot Solid-Phase-enhanced Sample Preparation (SP3) protocol^91,92^. To this end, lysates were added to Sera-Mag Beads (Thermo Scientific, #4515-2105-050250, 6515-2105-050250) in 10 µl 15% formic acid and 30 µl of ethanol. The binding of proteins was achieved by shaking for 15 min at room temperature. SDS was removed by 4 subsequent washes with 200 µl of 70% ethanol. Proteins were digested overnight at room temperature with 0.4 µg of sequencing grade modified trypsin (Promega, V5111) in 40 µl Hepes/NaOH, pH 8.4 in the presence of 1.25 mM TCEP and 5 mM chloroacetamide (Sigma-Aldrich, C0267). Beads were separated and washed with 10 µl of an aqueous solution of 2% DMSO, and the combined eluates were dried down.

Up to 10 µg of peptides per sample were labeled with TMT10plex™ reagents following established protocols^93^. Briefly, 0.8 mg of TMT reagent was dissolved in 45 µL of anhydrous acetonitrile, and 4 µL of this solution was added to each peptide sample. Labeling was carried out at room temperature for 1 h. The reaction was quenched by adding 4 µL of 5% hydroxylamine in water, followed by a 15-minute incubation at room temperature. Labeled peptides were pooled for multiplexing, desalted using an Oasis® HLB µElution Plate (Waters #186001828BA) according to the manufacturer’s instructions, and dried by vacuum centrifugation.

Offline high-pH reversed-phase fractionation was performed as described previously^94^, using an Agilent 1200 Infinity HPLC system equipped with a Gemini C18 analytical column (3 µm particle size, 110 Å pore size, 100 × 1.0 mm; Phenomenex) and a matching Gemini C18 SecurityGuard pre-column (4 × 2.0 mm; Phenomenex). Peptides were separated at a flow rate of 0.1 mL/min using mobile phase A (20 mM ammonium formate, pH 10.0) and mobile phase B (100% acetonitrile). The chromatographic gradient was as follows: 100% A for 2 min, linear increase to 35% B over 59 min, ramp to 85% B in 1 min, held at 85% B for 15 min, followed by re-equilibration at 100% A for 13 min. A total of 48 fractions were collected and subsequently pooled into 12 final fractions. Nanoflow LC-MS/MS analysis was performed on an UltiMate 3000 RSLCnano system (Thermo Fisher Scientific) coupled to an Orbitrap Fusion™ Lumos™ Tribrid™ mass spectrometer (Thermo Fisher Scientific). Peptides were initially loaded onto a µ-Precolumn C18 PepMap™ 100 trap cartridge (300 µm i.d. × 5 mm, 5 µm particles, 100 Å; Thermo) at 30 µL/min with 0.05% TFA in water for 6 min. Analytical separation was carried out using a nanoEase™ M/Z HSS T3 column (75 µm i.d. × 250 mm, 1.8 µm particles, 100 Å; Waters), equilibrated with solvent A (3% DMSO, 0.1% formic acid in water). Peptides were eluted at 0.3 µL/min using a gradient of solvent B (3% DMSO, 0.1% formic acid in acetonitrile) as follows: 2–8% B over 6 min, 8–25% over 99 min, 25–40% over 5 min, ramp to 85% in 0.1 min, held at 85% for 3.9 min, followed by re-equilibration to 2% B for 6 min. Mass spectrometry was performed using a Pico-Tip emitter (360 µm OD × 20 µm ID, 10 µm tip; CoAnn Technologies) with a spray voltage of 2.4 kV and a capillary temperature of 275 °C. Full MS scans were acquired in the Orbitrap at a resolution of 120,000 (at m/z 200), over a mass range of m/z 375–1,500, with an AGC target set to ‘standard’ and a maximum injection time of 50 ms. The instrument operated in data-dependent acquisition (DDA) mode. MS/MS spectra were acquired in the Orbitrap at a resolution of 30,000, with a maximum injection time of 94 ms and an AGC target of 200%. Precursors were fragmented using higher-energy collisional dissociation (HCD) at 36% normalized collision energy. A quadrupole isolation window of 0.7 m/z was used, with dynamic exclusion set to 60 s. Only precursor ions with charge states between 2 and 7 were selected for fragmentation.

Acquired data were analyzed using IsobarQuant and Mascot V2.4 (Matrix Science) using a reverse UniProt FASTA Mus musculus database (UP000000589), including common contaminants. The following modifications were considered: Carbamidomethyl (C, fixed), TMT10plex (K, fixed), Acetyl (N-term, variable), Oxidation (M, variable), and TMT10plex (N-term, variable). The mass error tolerance for full scan MS spectra was set to 10 ppm, and for MS/MS spectra to 0.02 Da. A maximum of 2 missed cleavages were allowed. A minimum of 2 unique peptides with a peptide length of at least seven amino acids and a false discovery rate below 0.01 were required on the peptide and protein level.

#### Data Processing

The data were processed using MaxQuant 2.1.3.0, with peptides identified from MS/MS spectra searched against the Uniprot Mouse Reference Proteome (UP000000589) using the built-in Andromeda search engine. Fixed modifications included cysteine carbamidomethylation, while variable modifications included methionine oxidation, glutamine/asparagine deamination, and protein N-terminal acetylation. In silico digests allowed for cleavages of arginine or lysine followed by any amino acid (trypsin/P), with up to two missed cleavages permitted. The false discovery rate (FDR) was set to 0.01 for peptides, proteins, and sites. Match between runs was enabled, and other parameters were used as preset in the software. Protein identification and intrasample quantification were performed using the iBAQ algorithm in MaxQuant.

### Transmission Electron Microscopy (TEM) of the Spleen

Fresh spleen samples, approximately 3 mm² in size, were fixed in 2.5% glutaraldehyde for 24 h at 4°C. Following fixation, the samples were washed in PBS and postfixed with 1% osmium tetroxide for 1 h. After rinsing with water, the samples were incubated in 1% aqueous uranyl acetate for 12 h at 4°C. The samples were then dehydrated at room temperature using increasing concentrations of ethanol, infiltrated with epoxy resin (Sigma-Aldrich, 45-359-1EA-F), and polymerized for 48 h at 60°C. The polymerized resin blocks were trimmed using a tissue processor (Leica EM TP) and sectioned into ultrathin slices (65 nm thick) with an ultramicrotome (EM UC7, Leica). These ultrathin sections were collected on nickel grids with 200 mesh (Agar Scientific, G2200N). The specimen grids were examined with a Tecnai T12 BioTwin transmission electron microscope (FEI, Hillsboro, OR, USA) equipped with a 16-megapixel TemCam-F416 camera (TVIPS GmbH) at the in-house Microscopy and Cytometry Facility.

### Preparation of Schematic Figures

The graphical abstract was created in BioRender. Mleczko-Sanecka, K. (2025) https://BioRender.com/y54j6da.

### Statistical Analysis

Female mice of the same age were randomly assigned to experimental groups. Samples derived from mice were collected and assessed/measured randomly, and harvested in a different order across individual experiments. The sample size, typically ranging from 5 to 9 mice per group or independent biological cell-based experiments, was determined based on power analysis, prior experience with similar experiments, and previously published studies in the field of RPM biology. Statistical analysis was conducted using GraphPad Prism (GraphPad Software, Version 9). Data are presented as mean ± SEM unless otherwise noted. Student’s t-test with Welch’s correction was employed when comparing the two groups. For comparisons involving more than two groups, one-way ANOVA was used, or two-way ANOVA when the simultaneous effects of two factors were being evaluated. Tukey’s test was implemented for post-hoc analysis following ANOVA. The following notation was used to indicate statistically significant differences between means: ns for non-significant (p > 0.05), * for p < 0.05, ** for p < 0.01, *** for p < 0.001, and **** for p < 0.0001.

## Data availability

Mass spectrometry proteomics data were deposited to the ProteomeXchange Consortium via the PRIDE partner repository with the dataset identifiers: Project accession: PXD064719 (access secured by a token: lKilHQArXhhB). RNA sequencing data are available in Gene Expression Omnibus GEO under the accession number GSE312948, and metadata were deposited to SRA (PRJNA1274426).

## Authorship Contributions

KMS, PKM, RM, KC and PS conceived and planned the experiments; PKM, RM, KC, PS, GZ, MN, AJ, and ML performed research; PKM, RM, KC, PS, GZ, KMS and ML analyzed and visualized data; EN, ZL and FG provided critical reagents/models, KC and KMS curated data; KMS, WP, and EN supervised the study; KC edited the manuscript; PKM, RM and KMS wrote the manuscript.

## Conflict of interest

EN is a shareholder and scientific advisor of Intrinsic LifeSciences and Silarus Therapeutics, and a consultant for Disc Medicine, Ionis Pharmaceuticals, Protagonist, GSK and Vifor. Other authors have declared that no conflict of interest exists.

## Acknowledgments

We thank the EMBL Proteomics Core Facility (EMBL, Heidelberg) for performing RPM proteomic profiling and data analysis, and the GeneCore team (EMBL, Heidelberg) for performing RNA sequencing. We thank Tara Arvedson (Amgen Inc., USA) for the anti-ferroportin antibody, Elizabeta Nemeth (UCLA, USA) for PR73, Florent Ginhoux for access to Ms4a3-Cre mice, and Aneta Suwińska for sharing UBI-GFP/BL6 mice, and Vania Manolova (CSL Vifor) for providing vamifeport. Many thanks to Agnieszka Popielska and Anna Kosson, and the staff of the Experimental Medicine Centre (Bialystok, Poland) and Mossakowski Medical Research Institute (Warsaw, Poland) for their technical support. We thank Sylwia Herman for her assistance with the histological analyses and Enrico Tatti for consulting on the MitoPlates profiling data. Mouse-based experiments and animal housing were supported by the Rodent Facility, EM, and Opera Phenix imaging were performed at the Microscopy Facility, and lentiviral DNA constructs were generated at the Genome Engineering Facility of IIMCB (IN-MOL-CELL Infrastructure) funded by the European Union – NextGenerationEU under the National Recovery and Resilience Plan. IN-MOL-CELL Infrastructure was also funded by the European Union under Horizon Europe (Project 101059801 - RACE) and by the RACE-PRIME project carried out within the IRAP programme of the Foundation for Polish Science, co-financed by the European Union under the European Funds for Smart Economy 2021-2027 (FENG). Pratik Kumar Mandal gratefully acknowledges funding from the National Science Centre PRELUDIUM 22 grant, which supported experiments involving lentiviral-mediated FPN overexpression (UMO-2023/49/N/NZ3/04232). This work was funded by the National Science Centre Sonata Bis grant (UMO-2020/38/E/NZ4/00511).

During the preparation of this work, the authors used ChatGPT 5.2 to assist with language editing in a limited number of instances. After using this tool, the authors reviewed and edited the content as needed and take full responsibility for the content of the published article.

## Notes

https://www.ncbi.nlm.nih.gov/geo/query/acc.cgi?acc=GSE312948

https://www.ebi.ac.uk/pride/archive/projects/PXD064719/privatereviewdataset

### Summary of Updates

Minor revisions were introduced in the references and the Discussion section, and Tables S1 and S2 have been added.

