## Supplementary Figures for "Iron deficiency drives metabolic adaptation of red pulp macrophages via ferroportin-SYK signaling and BCAA catabolism to enhance erythrophagocytosis"

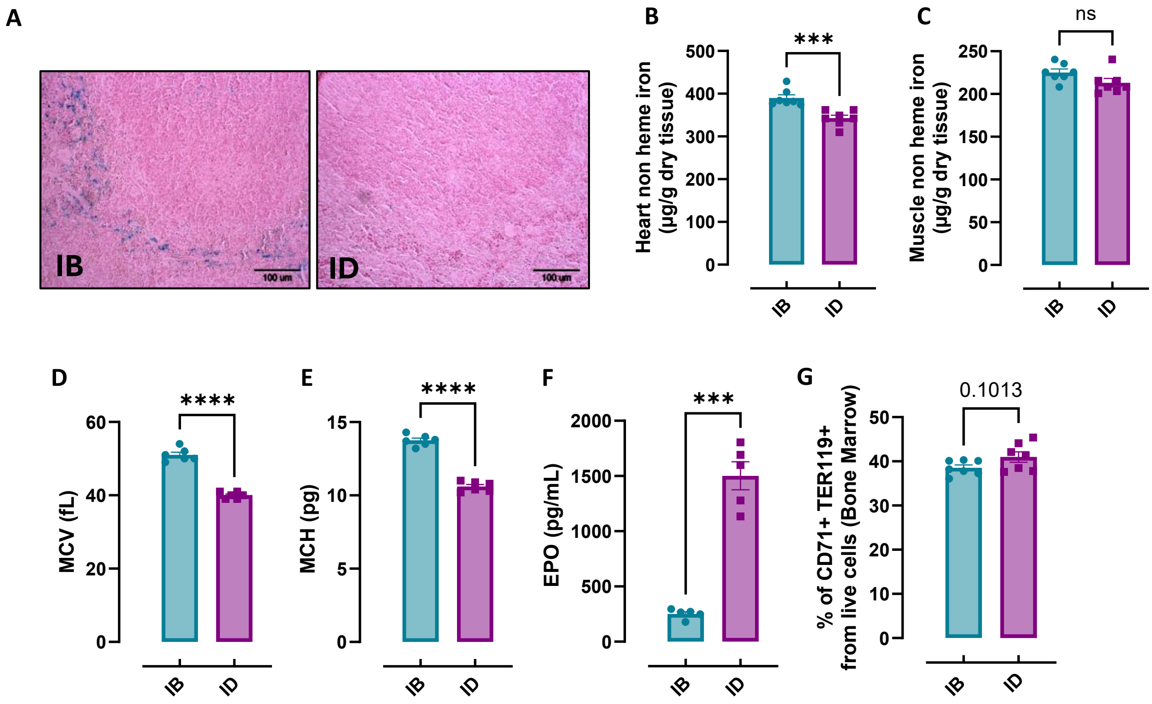


Figure S1. **Body iron indices and hematological parameters of nutritional iron-deficient mice.**
(A) Perls' Prussian Blue staining of the splenic red pulp in IB and ID mice. (B) Heart and (C) muscle non-heme iron content was determined in IB and ID mice. (D) Mean corpuscular volume (MCV), and (E) mean corpuscular hemoglobin (MCH) were determined in IB and ID mice. (F) EPO concentration in the serum of IB and ID mice was measured by Mouse Erythropoietin/EPO Quantikine ELISA Kit. (G) Shown is the percentage of erythroid progenitor cells (TER119+ CD71+) present in the bone marrow of IB and ID mice. Each dot represents one mouse. Data are represented as mean ± SEM. Welch's unpaired t-test determined statistical significance between the two groups. ns $p>0.05$, *** $p<0.001$ and **** $p<0.0001$.

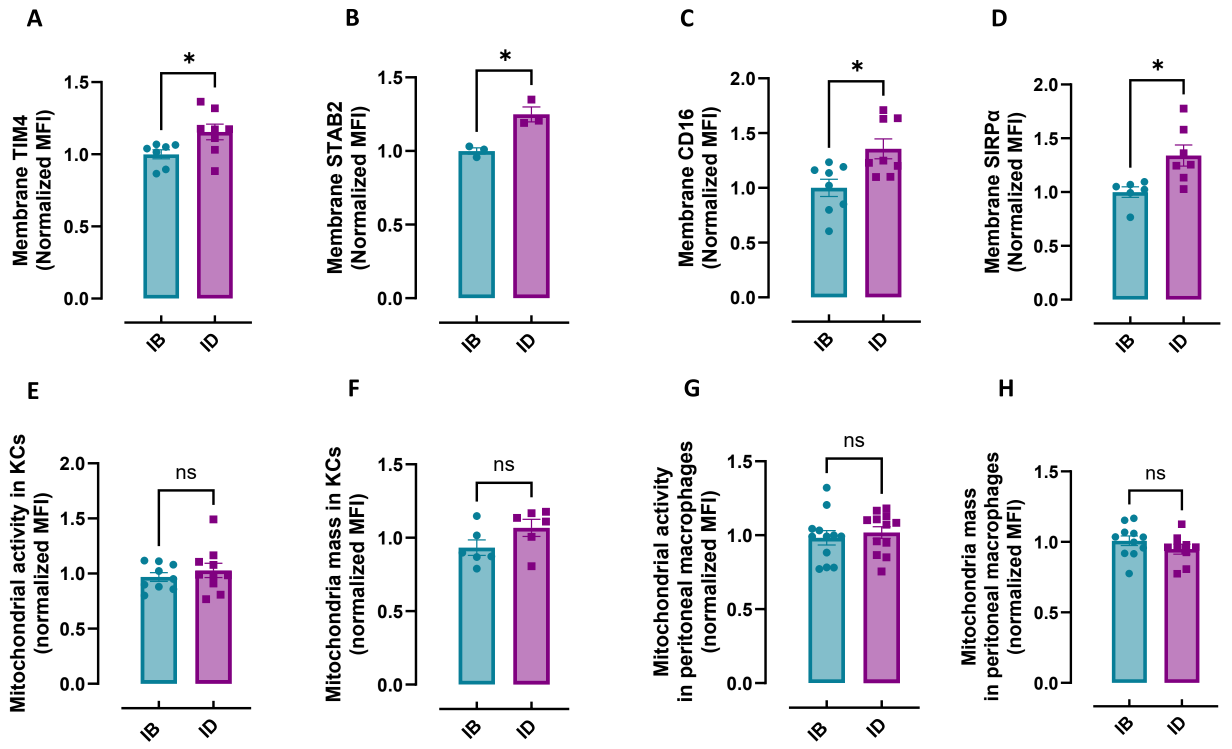


Figure S2.1: **Apoptotic cell receptors in RPMs and mitochondrial profiling of Kupffer cells and peritoneal macrophages.**

Cell membrane expression levels of (A) TIM4, (B) STAB2, (C) the Fc receptor CD16, and (D) SIRPα of RPMs in IB and ID mice were determined by using fluorescently labelled antibodies and flow cytometry. (E) Mitochondrial activity and (F) mitochondrial mass were determined in Kupffer cells (KCs) derived from IB and ID mice using TMRE probes and MitoTracker Green, respectively, with flow cytometry. (G) Mitochondrial activity and (H) mitochondrial mass were determined in peritoneal macrophages derived from IB and ID mice using TMRE probes and MitoTracker Green, respectively, with flow cytometry. Each dot represents one mouse. Data are represented as mean ± SEM. Welch's unpaired t-test determined statistical significance between the two groups. ns $p>0.05$, *$p<0.05$, **$p<0.01$, ***$p<0.001$ and ****$p<0.0001$.

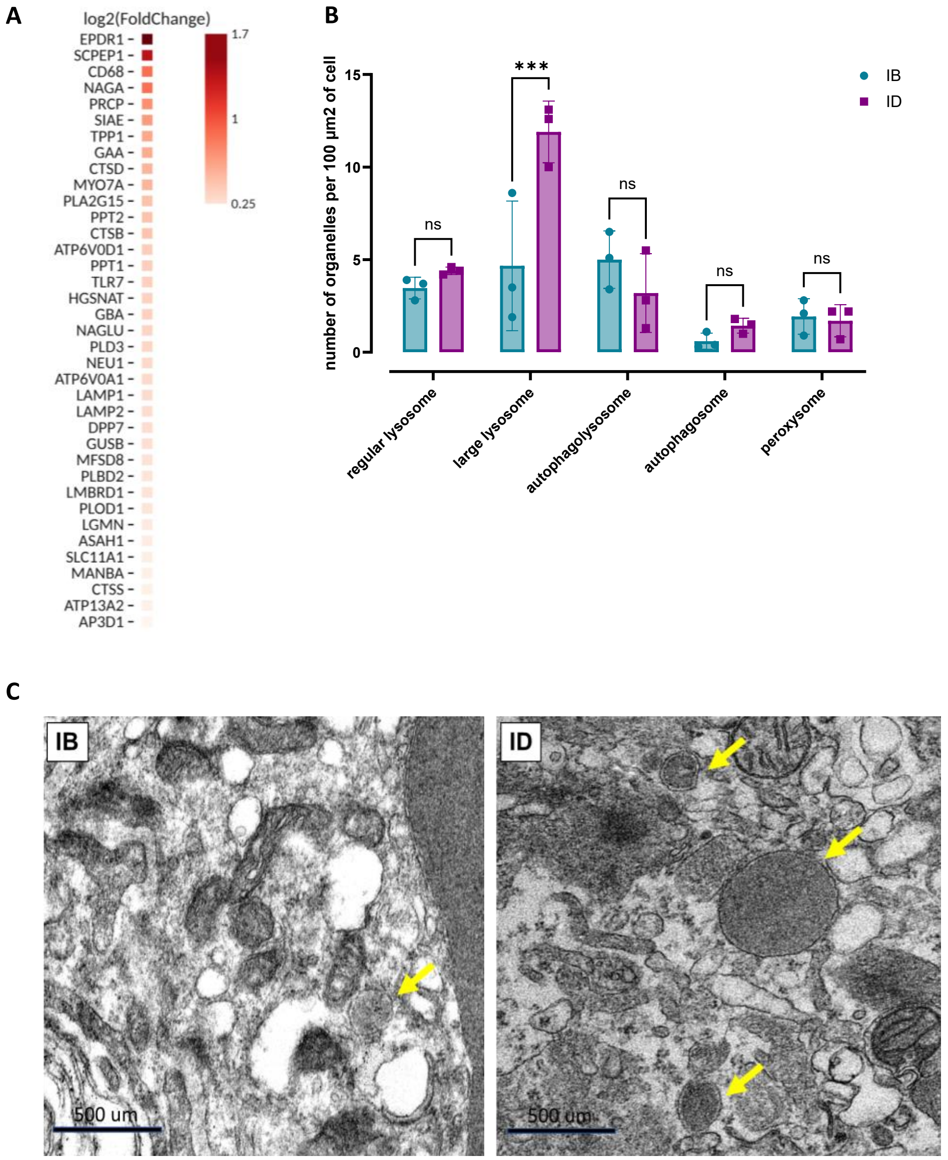


Figure S2.2. **Characterization of increased lysosomal function in RPMs of ID mice.**

(A) Heatmap showing $\log_2$ fold changes of lysosome-related proteins upregulated in RPMs from ID versus IB mice. (B) Quantification of regular lysosomes, large lysosomes, autophagolysosomes, autophagosomes, and peroxisomes per 100 µm² of cell area in RPMs from IB and ID mice. (C) Representative images of enlarged lysosomes in RPMs of IB and ID mice. Yellow arrows indicate enlarged lysosomes. Each dot represents one mouse. Data are represented as mean ± SEM. Welch's unpaired t-test determined statistical significance between the two groups. ns $p>0.05$, *$p<0.05$, **$p<0.01$, ***$p<0.001$ and ****$p<0.0001$.

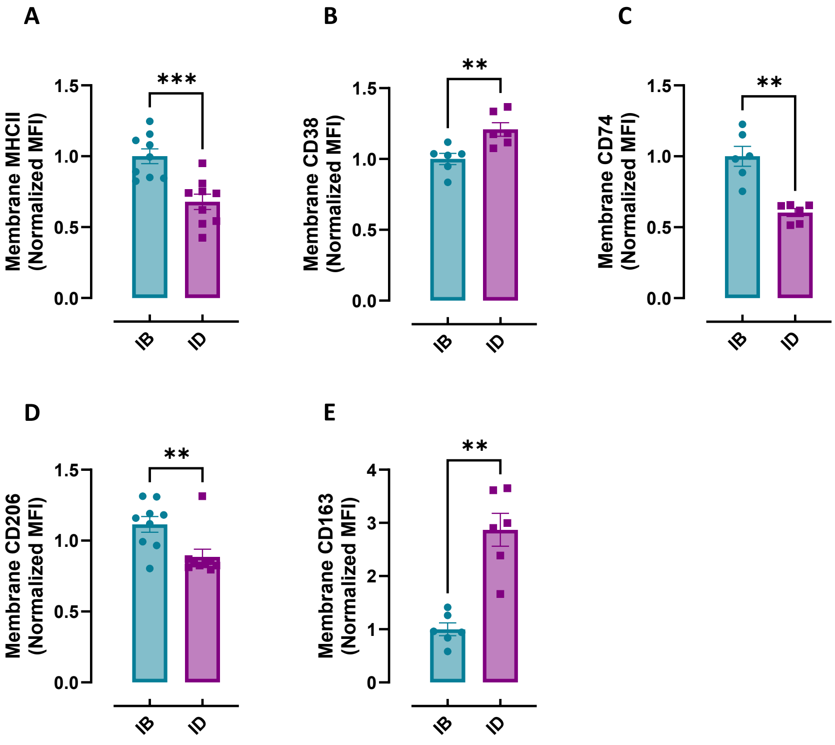


Figure S2.3. **Profiling of macrophage polarization markers in RPMs under ID conditions.**

Cell membrane expression levels of (A) MHCII, (B) CD38, (C) CD74, (D) CD206 and (E) CD163 of RPMs in IB and ID mice were determined by using fluorescently labelled antibodies and flow cytometry. Each dot represents one mouse. Data are represented as mean ± SEM. Welch's unpaired t-test determined statistical significance between the two groups. ns $p>0.05$, *$p<0.05$, **$p<0.01$, ***$p<0.001$ and ****$p<0.0001$.

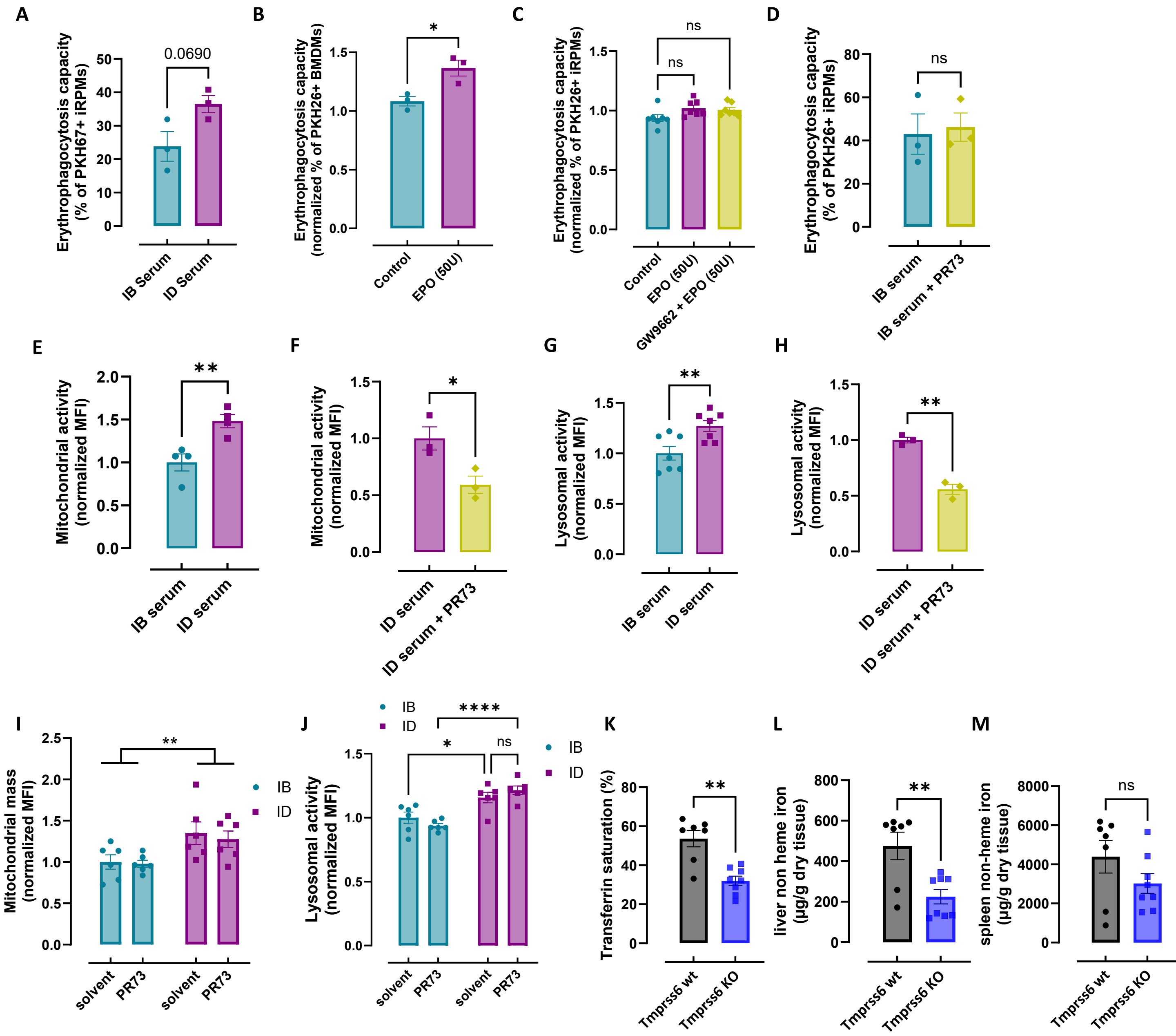


Figure S3: **The hepcidin-FPN axis regulates functional adaptations of RPMs during iron deficiency – extended data**

(A) The EP capacity in iRPMs treated with serum from IB or ID mice determined using flow cytometry by measuring the percentage of cells that phagocytosed transfused PKH26-labeled RBCs. (B) Normalized EP capacity of PKH26 sRBCs by cultured iRPMs treated with EPO determined using flow cytometry. (C) Normalized EP capacity of PKH26 sRBCs by cultured iRPMs with EPO (25 units, 24 h) treated alone or in combination with GW9662 (1µM, 25 h), a selective PPARγ antagonist determined using flow cytometry. (D) The EP capacity in iRPMs treated with serum from IB mice and serum from IB mice treated with the hepcidin agonist PR73 (50 nmol/mouse, 4 h) determined using flow cytometry by measuring the percentage of cells that phagocytosed transfused PKH26-labeled RBCs. (E) Mitochondrial activity was determined in iRPMs treated with serum from IB or ID mice using TMRE probe, with flow cytometry. (F) Mitochondrial activity was determined in iRPMs treated with serum from ID mice and serum from ID mice treated with the hepcidin agonist PR73 (50 nmol/mouse, 4 h) using TMRE probe determined using flow cytometry. (G) Lysosomal activity was determined in iRPMs treated with serum from IB or ID mice using the Lysosomal Intracellular Activity Assay Kit, with flow cytometry. (H) Lysosomal activity was determined in iRPMs treated with serum from ID mice and serum from ID mice treated with the hepcidin agonist PR73 (50 nmol/mouse, 4 h) using the Lysosomal Intracellular Activity Assay Kit determined using flow cytometry. (I) Mitochondrial mass was determined in RPMs from IB and ID mice using MitoTracker Green with flow cytometry after PR73 administration. (J) Flow cytometric assessment of lysosomal activity in RPMs from IB and ID mice using the Lysosomal Intracellular Activity Assay Kit following PR73 administration. (K) Plasma transferrin saturation was determined in Tmprss6 wt and Tmprss6 KO mice. (L) Liver and (M) spleen non-heme iron content was determined in wild type and Tmprss6 KO mice. Each dot represents one mouse or an independent cell-based experiment. Data are represented as mean ± SEM. Welch's unpaired t-test determined statistical significance in A-B, D, E-H and K-M, while two-way ANOVA with Tukey's Multiple Comparison tests was used in I-J. A one-way ANOVA test with Dunnett's or Tukey's Multiple Comparison tests was used in C. ns $p>0.05$, *$p<0.05$, **$p<0.01$, ***$p<0.001$ and ****$p<0.0001$.

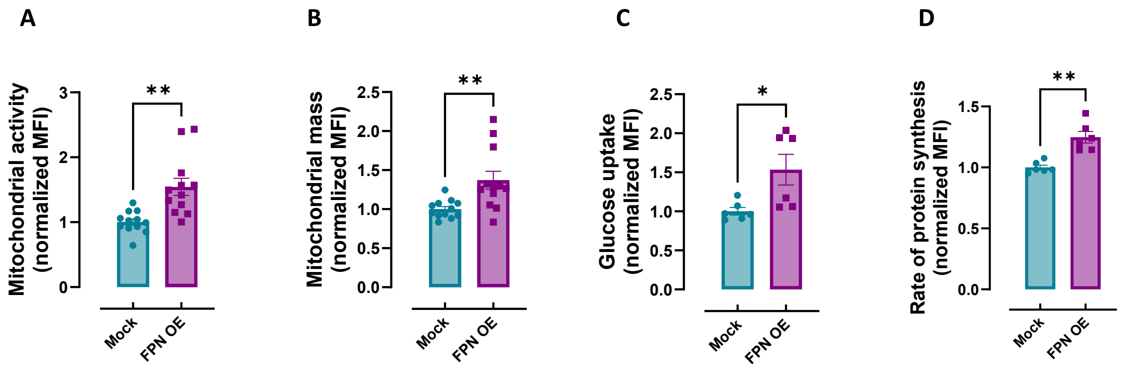


Figure S4. **Consequences of FPN overexpression on iRPMs functions in culture without sRBCs exposure.**

(A) Mitochondrial activity was determined in mock and FPN-iRPMs using TMRE probe, with flow cytometry. (B) Mitochondrial mass was determined in mock and FPN-iRPMs using MitoTracker Green, with flow cytometry. (C) Quantification of glucose uptake via flow cytometry in mock and FPN-iRPMs using a fluorescent glucose analog. (D) Flow cytometric detection of newly synthesized proteins in mock and FPN-iRPMs using Click-iT Plus OPP Alexa Fluor 488 Protein Synthesis Assay. Each dot represents an independent cell-based experiment. Data are represented as mean ± SEM. Welch's unpaired t-test determined statistical significance. ns $p>0.05$, *$p<0.05$ and **$p<0.01$.

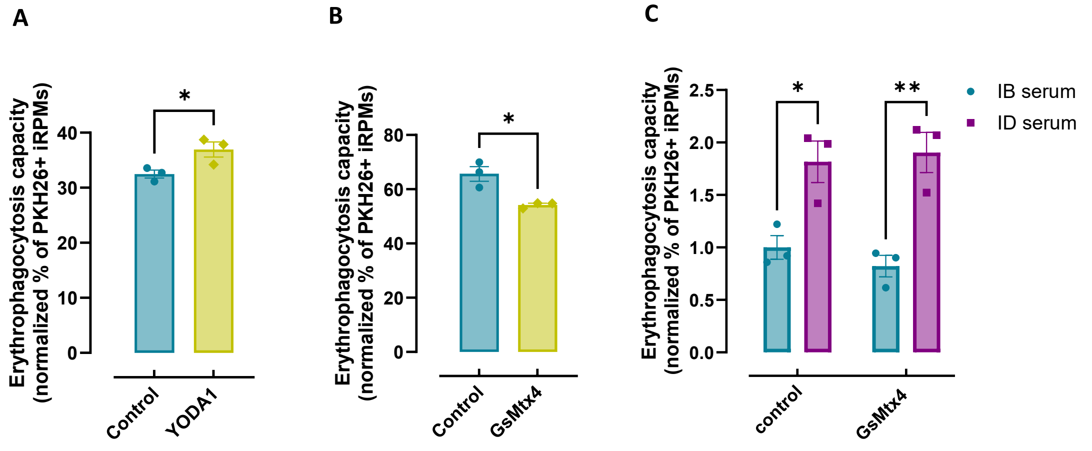


Figure S5.1. **The effects of PIEZO1 modulation in rewiring of RPMs in ID-like conditions.**

(A-B) EP capacity was assessed by flow cytometry as the percentage of iRPMs that internalized PKH26-labeled sRBCs, after treatment with (A) the PIEZO1 activator YODA1 (10 µM, 1h) or (B) the PIEZO1 inhibitor GsMTx4 (10 µM, 1h). (C) EP capacity was measured in iRPMs cultured with IB and ID serum by flow cytometry as the percentage of cells that phagocytosed PKH26-labeled sRBCs, upon treatment with GsMTx4. Each dot represents an independent cell-based experiment. Data are represented as mean ± SEM. Welch's unpaired t-test determined statistical significance in A-B. Two-way ANOVA with Tukey's Multiple Comparison tests was used in C. ns $p>0.05$, *$p<0.05$, **$p<0.01$,.

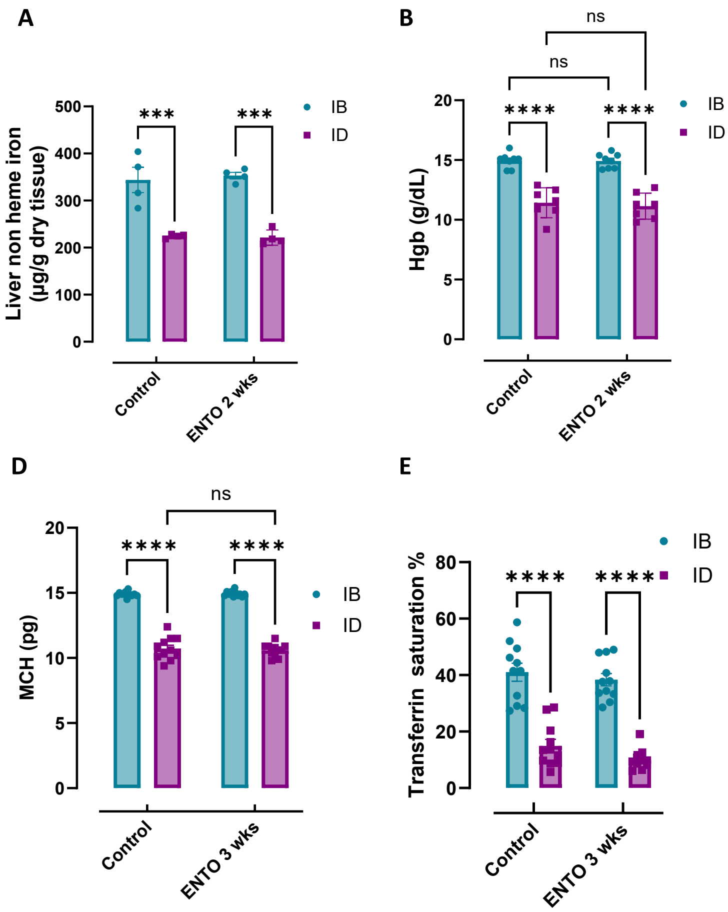


Figure S5.2. **SYK kinase inhibition disrupts the adaptation of RPMs to ID conditions in mice – extended data.**

(A) Liver non-heme iron content measured in IB and ID mice following 2 weeks of dietary intervention with or without ENTO jelly supplementation. (B) Blood hemoglobin (Hgb) levels in IB and ID mice after 2 weeks of ENTO jelly or control treatment. (C) Mean corpuscular hemoglobin (MCH) levels in IB and ID mice following 2 weeks of supplementation with ENTO jelly. (D) Mean corpuscular hemoglobin (MCH) levels in IB and ID mice following 3 weeks of supplementation with ENTO jelly. (E) Plasma transferrin saturation levels were measured in IB and ID mice after 3 weeks of supplementation with ENTO jelly. Each dot represents one mouse. Data are represented as mean ± SEM. Two-way ANOVA with Tukey's Multiple Comparison tests was used to determine statistical significance. ns $p>0.05$, *$p<0.05$, **$p<0.01$, ***$p<0.001$ and ****$p<0.0001$.

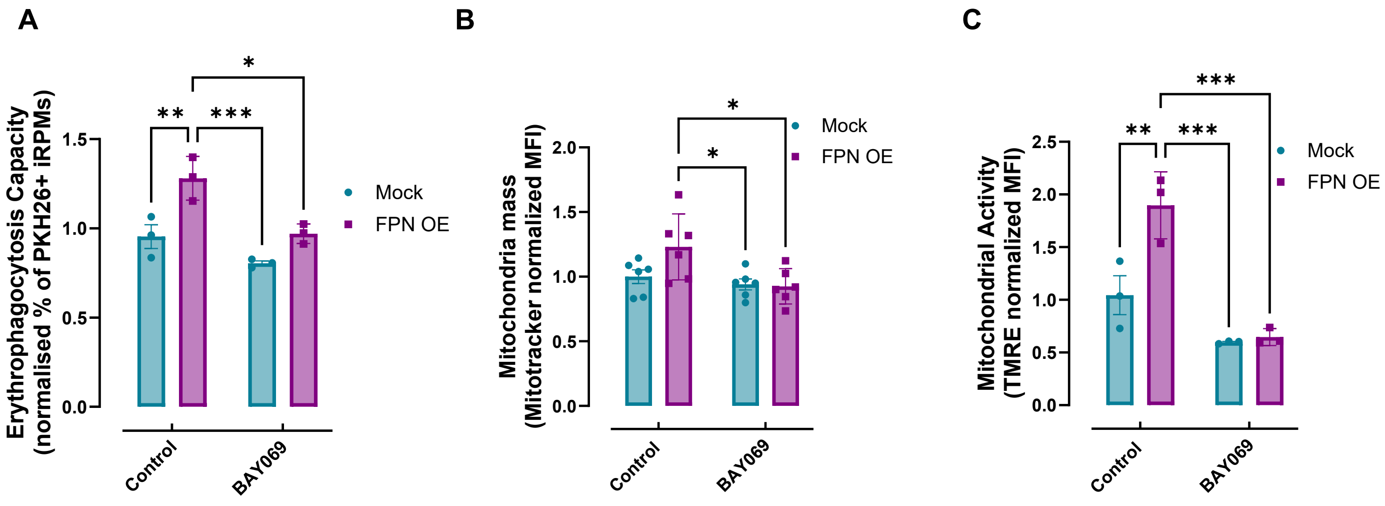


Figure S6.1. **Enhanced phagocytic and metabolic function of ID-like RPMs are reversed by BCAT inhibitor BAY069.**

(A) Normalized EP capacity of mock and FPN OE iRPMs upon treatment with BAY069 (10 µM, 15 h) measured by uptake of PKH26-labeled sRBCs. (B) Mitochondrial mass was determined in mock and FPN OE iRPMs upon treatment with BAY069 using MitoTracker Green, with flow cytometry (C) Mitochondrial activity was determined in mock and FPN OE iRPMs upon treatment with BAY069 using TMRE probe, with flow cytometry. Each dot represents an independent cell-based experiment. Data are represented as mean ± SEM. Two-way ANOVA with Tukey's Multiple Comparison tests was used to determine statistical significance. ns $p>0.05$, $*p<0.05$, $**p<0.01$, $***p<0.001$ and $****p<0.0001$.

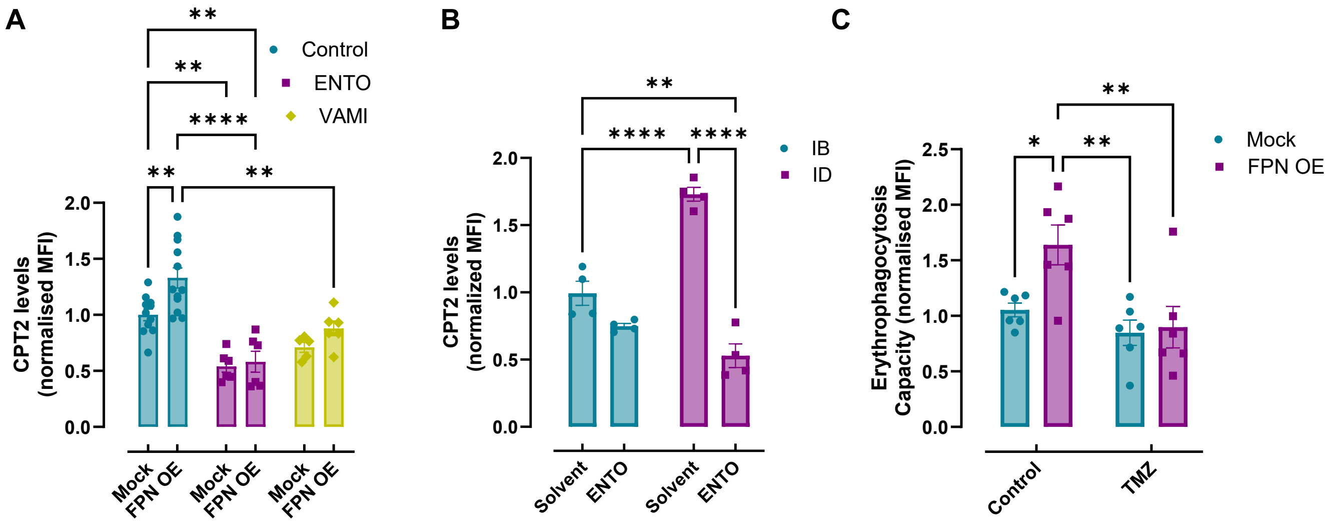


Figure S6.2. **Enhanced β-oxidation supports RPM adaptation to ID conditions.**

(A) Intracellular CPT2 levels in mock and FPN OE iRPMs upon treatment with Entospletinib (ENTO) (10 µM, 1 h) and vamifeport (VAMI) (20 µM, 15 h), determined using flow cytometry. (B) Intracellular CPT2 levels of RPMs from IB and ID mice supplemented with ENTO via jelly feeding (3 weeks), determined using flow cytometry. (C) Normalized EP capacity towards PKH26-sRBCs by mock and FPN OE iRPMs upon Trimetazidine (TMZ) treatment (10 µM, 15 h). Each dot represents one mouse or an independent cell-based experiment. Data are represented as mean ± SEM. Two-way ANOVA with Tukey's Multiple Comparison tests was used to determine statistical significance. ns $p>0.05$, *$p<0.05$, **$p<0.01$, ***$p<0.001$ and ****$p<0.0001$.

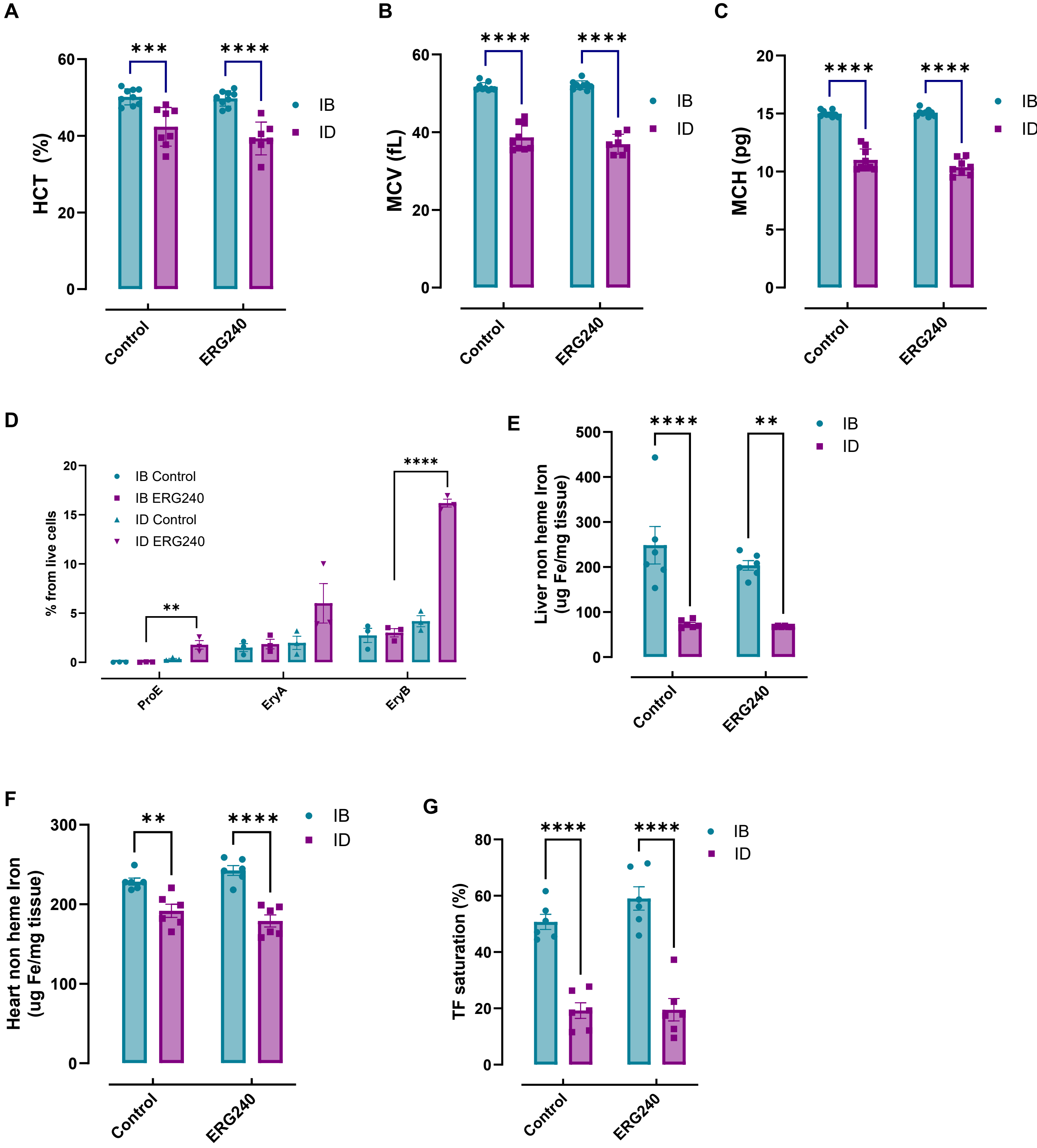


Figure S7. **In vivo inhibition of BCAA catabolism alters erythrophagocytic capacity and metabolic remodeling of RPMs in ID mice – extended data.**

(A) Hematocrit (HCT), (B) Mean corpuscular volume (MCV) and (C) mean corpuscular hemoglobin (MCH) were determined in IB and ID mice following 3 weeks of dietary ERG240 supplementation. (D) Proerythroblasts, EryA, and EryB erythroblast representations in the spleens of IB and ID mice supplemented with ERG240. (E) Liver and (F) heart non-heme iron content measured in IB and ID mice following 3 weeks of dietary ERG240 supplementation. (G) Plasma transferrin saturation was determined in IB and ID mice supplemented with ERG240. Each dot represents one mouse. Data are represented as mean ± SEM. Two-way ANOVA with Tukey's Multiple Comparison tests was used to determine statistical significance. ns $p>0.05$, *$p<0.05$, **$p<0.01$, ***$p<0.001$ and ****$p<0.0001$.
